# 3D microenvironment stiffness regulates tumor spheroid growth and mechanics via p21 and ROCK

**DOI:** 10.1101/586784

**Authors:** Anna V. Taubenberger, Salvatore Girardo, Nicole Träber, Elisabeth Fischer-Friedrich, Martin Kräter, Katrin Wagner, Thomas Kurth, Isabel Richter, Barbara Haller, Marcus Binner, Dominik Hahn, Uwe Freudenberg, Carsten Werner, Jochen Guck

**Author notes:** **Correspondence to:** Anna V. Taubenberger, TU Dresden Center for Molecular and Cellular Bioengineering (CMCB), Mailto.

## Abstract

Mechanical properties of cancer cells and their microenvironment contribute to breast cancer progression. While mechanosensing has been extensively studied using two-dimensional (2D) substrates, much less is known about it in a physiologically more relevant 3D context. Here we demonstrate that breast cancer tumor spheroids, growing in 3D polyethylene glycol-heparin hydrogels, are sensitive to their environment stiffness. During tumor spheroid growth, compressive stresses of up to 2 kPa built up, as quantitated using elastic polymer beads as stress sensors. Atomic force microscopy (AFM) revealed that tumor spheroid stiffness increased with hydrogel stiffness. Also, constituent cell stiffness increased in a ROCK- and F-actin-dependent manner. Increased hydrogel stiffness correlated with attenuated tumor spheroid growth, a higher proportion of cells in G0/G1 phase and elevated levels of the cyclin-dependent kinase inhibitor p21. Drug-mediated ROCK inhibition reversed not only cell stiffening upon culture in stiff hydrogels but also increased tumor spheroid growth. Taken together, we reveal here a mechanism by which the growth of a tumor spheroid can be regulated via cytoskeleton rearrangements in response to its mechanoenvironment. Thus, our findings contribute to a better understanding of how cancer cells react to compressive stress when growing under confinement in stiff environments and provide the basis for a more in-depth exploration of the underlying mechanosensory response.

## Introduction

Breast cancer is the most commonly diagnosed cancer in women worldwide, with a drastically decreased five-year survival rate after disease progression to an advanced stage.^[1]^ Multiple factors are known to contribute to tumor progression. Among them, ‘loss of mechanical reciprocity’ describes a progressive loss of tissue homeostasis and perturbations in tissue architecture,^[2,3]^ leading to an altered mechanical phenotype of tumor cells themselves and also of the surrounding tumor stroma.^[4]^ Clinically, a stiffened tumor stroma is reflected by increased mammographic density, which actually presents a strong risk factor of invasive breast cancer.^[5]^ The increase in stromal tissue stiffness has been attributed to excessive accumulation of extracellular matrix (ECM) molecules, such as collagen type I, tenascin-C and fibrin due to the action of cancer-associated fibroblasts.^[6][7]^ Nevertheless, only a few studies have compared the mechanical phenotype of normal and malignant mammary tissues at the cellular level. ^[8,9]^ Plodinec et al. detected a more heterogenous mechanical signature of malignant tissue in patient biopsies compared to normal tissue.^[8]^ Concurrently, using a mouse model of breast cancer, Lopez et al. reported an increase in tissue stiffness of malignant tissues over normal and pre-malignant epithelium.^[9]^ It is well established that cancer cells are sensitive to such changes in the mechanical properties of their environment. On the one hand, a stiffened stromal ECM can increase cytoskeletal tension in breast cancer cells, activate integrins and thereby modulate signaling pathways regulating cell differentiation, proliferation, invasion and treatment response.^[2,4]^ On the other hand, the expanding tumor pushes against the surrounding stromal compartment, which causes not only its deformation and structural rearrangement, but also results in compressive stress acting on the tumor cells.^[10,11]^ A stiffened microenvironment is expected to amplify this stress even more. Studies that investigated the effect of compressive stress on the growth of different cancer cells *in vitro*, either as spheroid cultures in inert agarose gels or free-floating spheroids exposed to increased osmotic pressure,^[12,13,14]^ typically reported reduced spheroid sizes. Furthermore, the effect of hydrostatic pressure on 2D breast cancer cell cultures was tested and found to induce a migratory/invasive phenotype.^[15]^

It is well acknowledged by now that 3D cultures represent a physiologically more relevant culture system for breast cancer cells compared to 2D cultures.^[16,17]^ 3D cultures show marked differences from 2D cultures with respect to cell morphology, gene expression, signal transduction, proliferation, migration and drug resistance.^[18,19]^ Different approaches exist to generate 3D *in vitro* models. For instance, tumor spheroids generated by hanging droplets assemble from multiple cells, but do not fully replicate the ECM microenvironment of native tissues. Embedding cells into 3D matrices assembled from different biomaterials alters solute diffusion, binding of growth factors and other effector proteins, thereby creating more realistic tissue-scale concentration gradients.^[16]^ While biomaterials of natural origin, such as collagen type I, provide a fibrillar ECM microenvironments, they do not allow for independent tuning of biochemical factors/ligand density and mechanical properties. In contrast, synthetic hydrogels permit independent modification of biochemical (e.g. matrix metalloproteinase (MMP) cleavage, ligand binding sites) and mechanical features of the created tumor microenvironment.^[19–21]^

In order to systematically study the effect of a stiffened microenvironment on tumor spheroids, we cultured here MCF-7 breast cancer spheroids in PEG-heparin based 3D microenvironments. This 3D model incorporated important aspects of physiological tumor microenvironments such as matrix degradability, while its mechanical properties were tunable without affecting ligand density.^[22]^ We quantitatively showed using cell-scale stress sensors, that stiff hydrogels caused increased compressive stress on the growing spheroid. Increased compression was accompanied by elevated nuclear p21 levels, delayed cell cycle progression and attenuated spheroid growth. Reduced spheroid growth in stiff microenvironments coincided with a ROCK dependent increase in the stiffness of constituent cells. Interfering with ROCK signaling not only reversed this cell stiffening response but also increased spheroid growth. Together, we describe here a mechanism by which tumor spheroid growth can be regulated via cytoskeleton alterations in response to the tumor spheroids’ 3D mechanoenvironment.

## Results

### Engineering 3D tumor microenvironments of defined mechanical properties

To generate 3D cultures mimicking avascular tumors, MCF-7 breast cancer cells were embedded into MMP-degradable PEG-heparin hydrogels as previously described (Figure 1A).^[22,23]^ During the 14-day culture period, multicellular spheroids formed from the single cell stage and reached diameters of up to 150 µm (Figure 1B). To recreate a stiffness range encompassing the reported stiffness of breast cancer tissue,^[8]^ the PEG-heparin ratio (γ) was adjusted to values of 0.75 (compliant), 1.0 (intermediate) and 1.5 (stiff), yielding hydrogels with Young’s moduli ranging from approximately 2 to 20 kPa (Figure 1C). AFM indentation tests showed that the hydrogels were predominantly elastic and maintained their global mechanical properties over the maximum culture period of 14 days (Figure S1). Since the amount of the PEG component was altered, while the bioactive heparin component was maintained constant, this allowed for uncoupling of mechanical and biochemical cues (Figure1D). MCF-7 cells formed spheroids over the entire gel stiffness range (Figure1E) but appeared smaller in the stiff hydrogels. Thus, we next set out to characterize their morphology in more detail.

**Figure 1.**
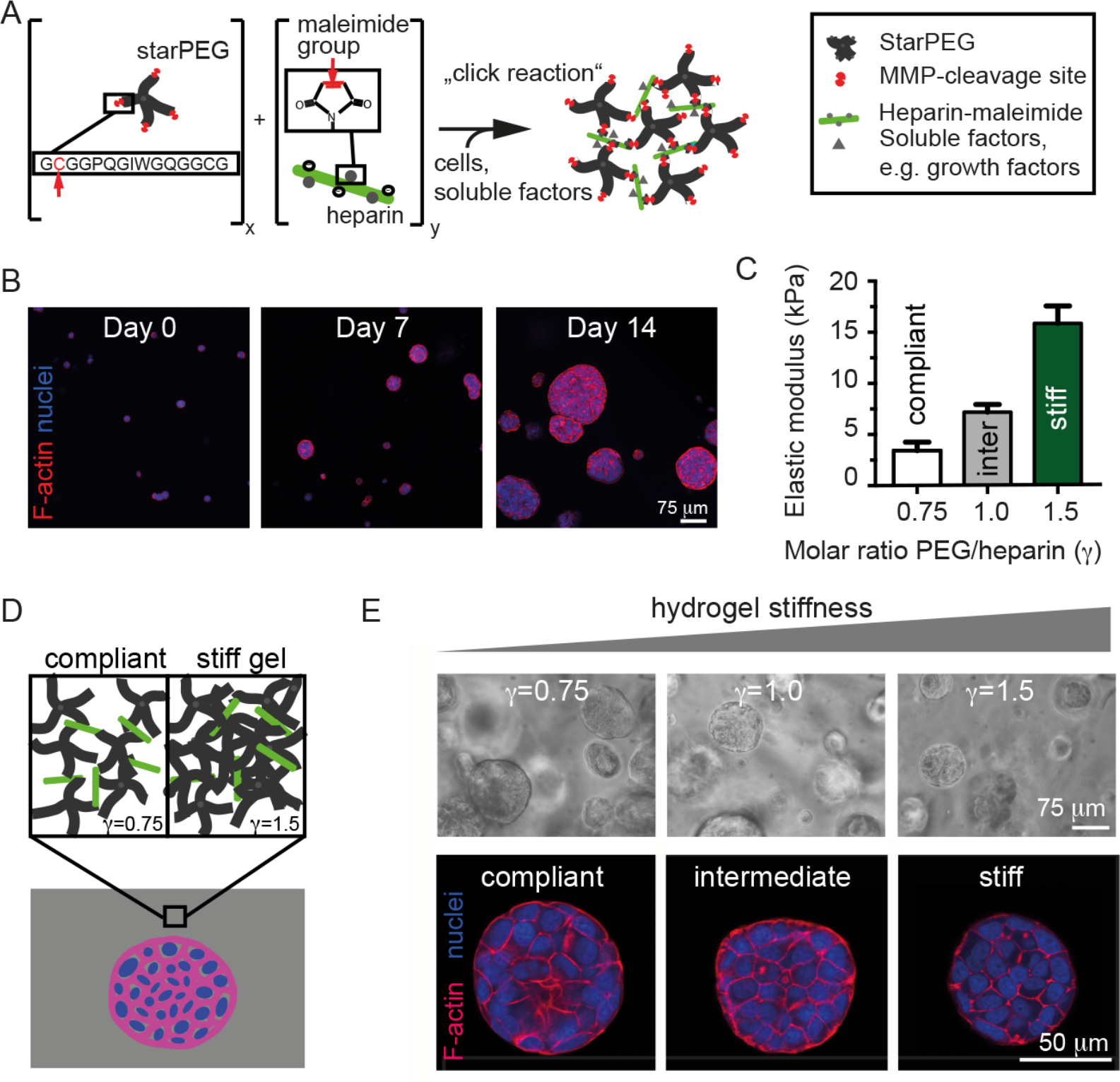
MCF-7 tumor spheroid formation in hydrogels of defined mechanical properties. **A.** Schematic illustration showing the hydrogel chemistry. Cysteine residues within the four-arm PEG and maleimide-modified heparin (red arrows) are covalently coupled by click chemistry, yielding a crosslinked hydrogel matrix around single MCF-7 cells suspended in the precursor mix. The PEG precursors contain MMP-cleavage sites and thereby allow for local degradation of the hydrogel matrix by cells **B.** Representative confocal images of MCF-7 cells/spheroids showing F-actin (red) and nuclei (blue) at different culture days. Cells embedded into the hydrogel mixture grow from the single cell stage into multicellular tumor spheroids over the culture period of 14 days. **C.** Characterization of the mechanical properties of hydrogels by AFM indentation. Depending on the molar ratio of PEG/heparin-maleimide, hydrogels of different stiffness (Young’s modulus) can be generated. Data are presented as mean ± standard error of the mean (SEM). **D.** By adjusting the molar ratio of PEG whilst maintaining the heparin-maleimide amount constant, hydrogel stiffness can be tuned independently of ligand density. **E.** Representative phase contrast (above) and confocal images of F-actin (red) and nuclei (blue) stained spheroids (below) growing in hydrogels of varying stiffness.

### Spheroids are smaller and more compact when grown in stiff microenvironments

To test how microenvironment stiffness affects spheroid morphology and size, 14-day-old spheroids were either analyzed *in situ* or after release from the hydrogel by collagenase-mediated matrix degradation (Figure 2A). We observed a significant reduction in spheroid size with increasing hydrogel stiffness (about 30% decrease on average for stiff versus compliant) (Figure 2B, C), while spheroid shapes were comparable over the entire stiffness range (Figure 2D, E). Consistent with spheroid size decrease, less cells were typically found per cross sections of spheroids grown in stiffer hydrogels (Figure 2F, Figure S2). Normalization to the respective cross-sectional area revealed a ~18% higher density of nuclei in stiff gels (Figure 2G). This compaction was observed both in cryosections and confocal images of intact hydrogel cultures (Figure 1E). The same phenomenon of reduced spheroid size with increasing hydrogel stiffness was also found for two other cell lines, T47D and LNCaP cells (Figure S3). Confocal images of MCF-7 spheroids further revealed occasional lumen formation (Figure S2), which was mostly seen in compliant gels (22%), and less frequently in stiffer gels (10 and 6% for intermediate and stiff, respectively, Figure 2I). Transmission electron microscopy (TEM) images also revealed lumina and vesicles indicative of secretory activity, and confirmed that spheroids were vital with no necrotic cores (Figure S4). Moreover, in compliant gels cells typically extended multiple filopodia towards the gel matrix, while in stiff gels the cell membrane at the external spheroid surface appeared smoother with less filopodia. Cells appeared pushing against the gel structure at some locations around the spheroid surface, as suggested by the locally increased gel density and gel deformation (Figure 2H, arrows).

**Figure 2.**
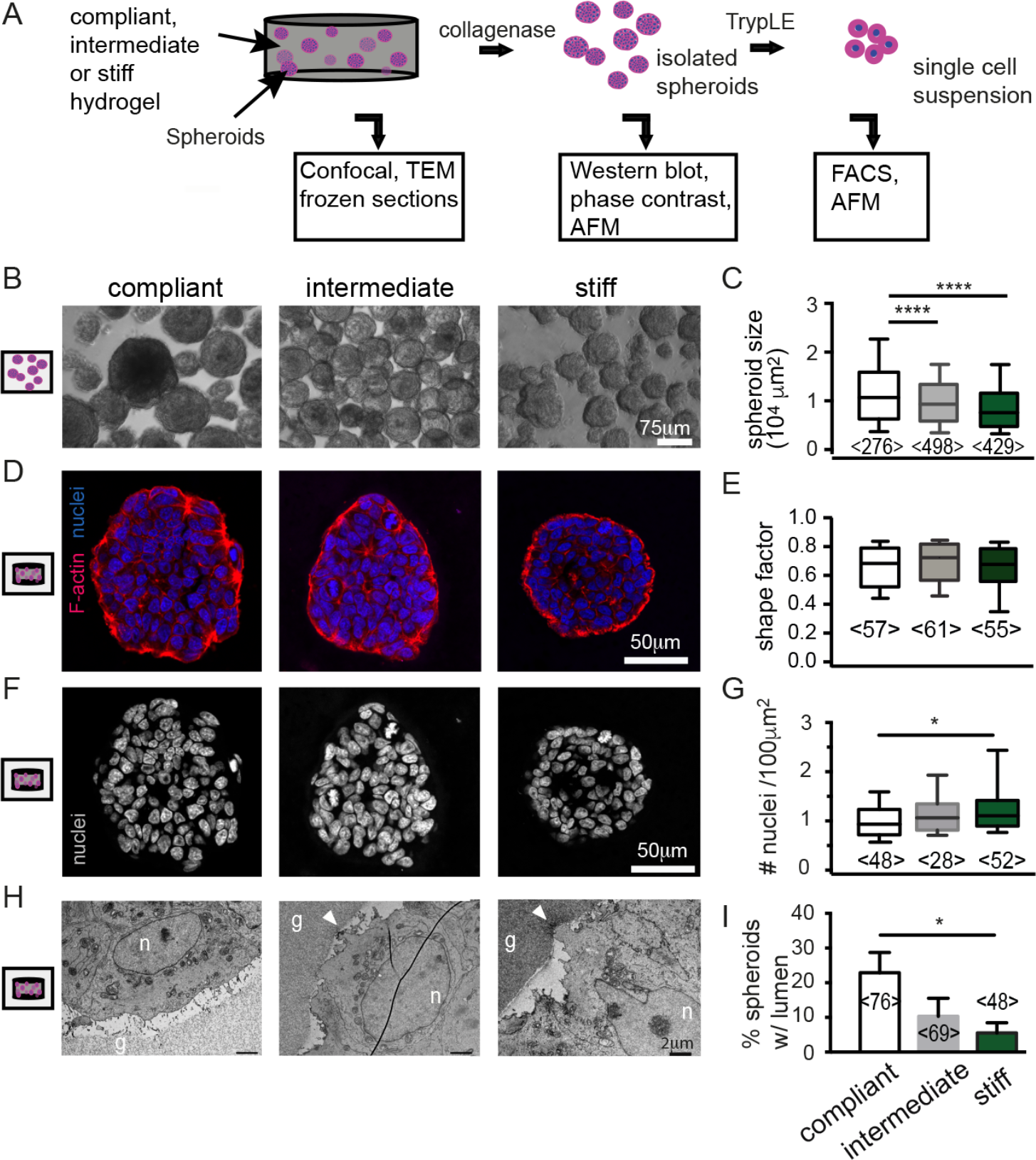
Morphological characterization of tumor spheroids grown in microenvironments of varying stiffness. **A.** Schematic illustration showing the processing of hydrogels to characterize intact cultures, isolated spheroids or single cells. **B.** Phase contrast images of spheroids released from the hydrogels. **C.** Quantification of spheroid size (diameter; >3 independent experiments). **D.** Confocal images of cryosections stained for F-actin (red) and nuclei (blue). **E.** Shape factors determined from confocal *in situ* images of F-actin stained spheroids (e.g. Figure 1E). **F.** DAPI channel of D. **G.** Using the DAPI channel, the number of nuclei was counted in confocal images of cryosections (e.g. Figure 2F) and normalized to the respective tumor spheroid cross sectional area (measured in FIJI using the F-actin signal). Data in **C,E,G** are presented as Box-Whisker plots, where boxes mark the 25, 50 and 75 percentile, and whiskers mark the 10 and 90 percentiles. **H.** TEM images of the spheroid/hydrogel boundary. Nuclei are marked with “n”, hydrogel matrix with “g”. Arrows highlight spots of gel deformation. **I.** Percentage of spheroids with lumen as determined in confocal images of F-actin/nuclei stained 3D cultures *in situ*. Data are presented as mean ± SEM. Percentages were calculated per hydrogel (at least n=7). Numbers in brackets indicate the number of analyzed tumor spheroids. Where indicated, datasets were compared using an ANOVA test with a Tukey’s multiple comparison test. **** denotes p-values < 0.001, * p < 0.05.

### MCF-7 tumor spheroids experience compressive stress in stiff microenvironments

To test whether tumor spheroids were exposed to compressive stress in stiffer hydrogel cultures, monodisperse cell-sized elastic polyacrylamide beads of controlled mechanical properties were employed as stress sensors.^[24]^ The stress sensors were mixed together with single cells into the PEG-heparin hydrogels during gel formation, and 3D cultures were maintained for 14 days to allow for tumor spheroid growth (Figure 3A). Confocal images showed that beads were considerably deformed when in the proximity of the spheroids in stiff, but not compliant hydrogels (Figure 3B). Radial stresses, causing this bead deformation, were estimated using the beads’ elastic modulus (4 kPa) and aspect ratio, and plotted against the radial distance from the spheroid edge (Figure 3A, C). The radial stress exponentially decayed within 40-50 μm distance from the spheroid, which was approximated by an exponential function. Beads close to the edge of large spheroids (>100 μm diameter) encountered radial stresses of approximately 2 kPa in stiff hydrogels. To test if compressive stress had been released in compliant gels by matrix degradation, beads and cells were also embedded into non-degradable PEG-heparin gels lacking MMP cleavage sites. Under these conditions, bead deformation was seen in both, compliant and stiff hydrogels, and stresses were comparable to degradable hydrogels (Figure 3D). However, tumor spheroids reached considerably smaller sizes in non-degradable than in degradable hydrogels (Figure 2, Figure S5). In non-degradable hydrogels, tumor spheroids also exhibited a more rounded morphology and smoother edges than in degradable gels (Figure S5). Together, these findings suggested that in stiff hydrogels a higher level of compressive stress is built up around spheroids. We thus questioned how the cells mechanically balanced increased compressive forces and set out to quantitatively compare the mechanical properties of spheroids/cells cultured in compliant and stiff hydrogels.

**Figure 3.**
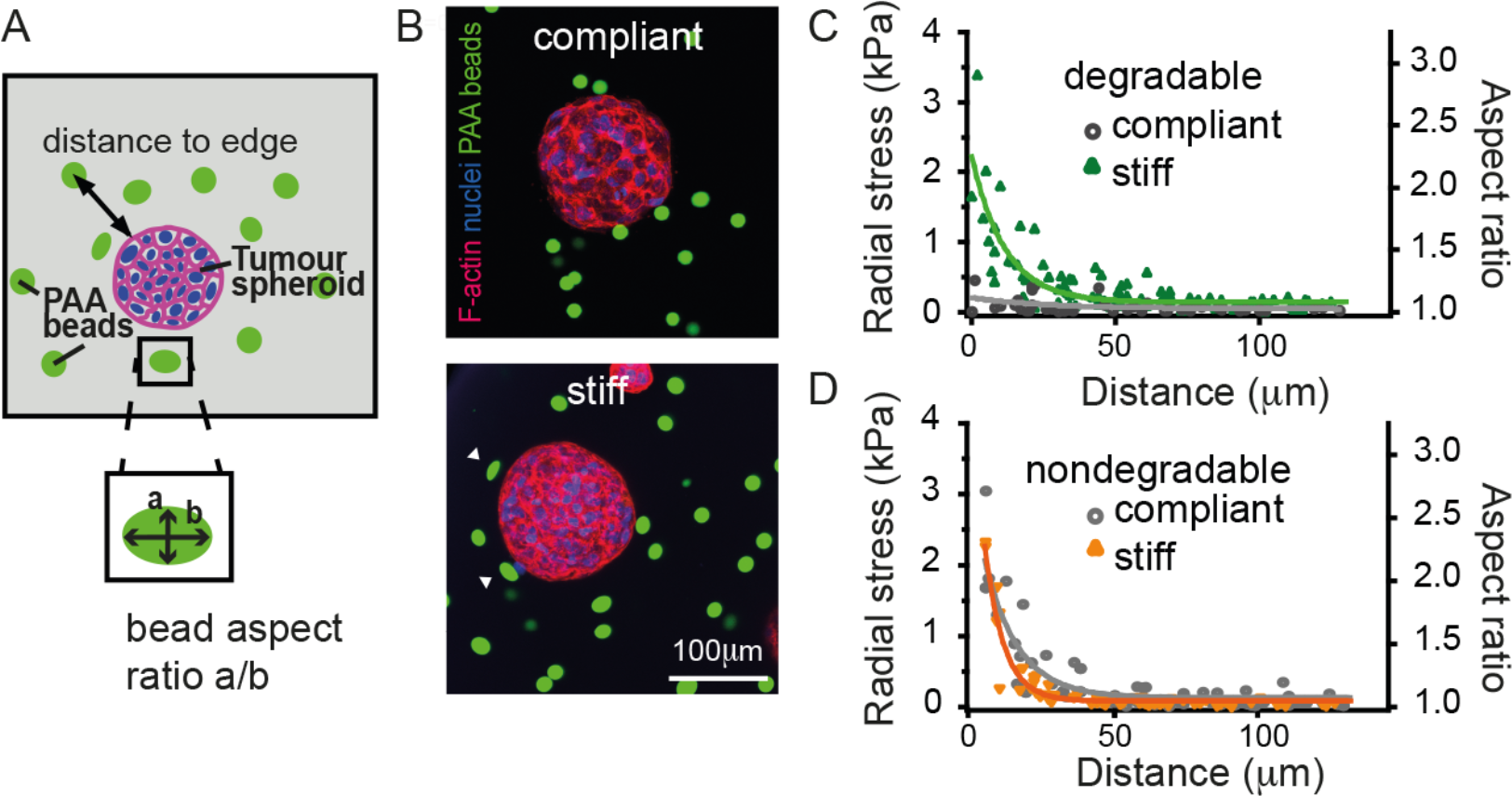
Quantification of growth-induced compressive stress acting on tumor spheroids. **A.** To test if compressive stress builds up during tumor spheroid growth, fluorescently (Cy3-) conjugated polyacrylamide beads with a diameter of 15 µm were employed as stress sensors. 14-day tumor spheroid cultures fixed and stained for F-actin (red) and nuclei (blue). Bead aspect ratio, spheroid diameter, and distance to spheroid edge were measured using Fiji and Zen software, respectively. **B.** Representative confocal images of stained spheroids. White arrowheads highlight deformed beads. **C.** Aspect ratio of beads and corresponding radial stress plotted over the distance from the tumor spheroid edge for degradable hydrogels. Single data points correspond to single beads, pooled from different images/tumor spheroids of comparable diameter (>100 µm). **D.** Aspect ratio and corresponding radial stress plotted over the distance from the spheroid edge for non-degradable gels (all spheroid diameters pooled). Single data points correspond to single beads. Data points in C and D were fit with an exponential function.

### Cells have increased stiffness in stiff microenvironments

The spheroids’ mechanical properties were probed at different scales. Spheroids were firstly isolated from hydrogels by collagenase treatment and probed on their surface by AFM indentation. Significantly increased apparent Young’s moduli were obtained for tumor spheroids grown in stiff hydrogels (Figure 4A). Since the probe size (10 µm bead) was smaller than the cell diameter, this effect was largely attributed to increased stiffness of cells lining the spheroid surface. Additionally, SEM images did not indicate extensive ECM deposition on the spheroids’ surface (Figure S6). AFM indentation on individual cells after tumor spheroid dissociation also revealed higher apparent Young’s Moduli for cells originating from stiff hydrogel cultures. Since cells were only indented to a maximum of 0.5 µm, this pointed to an increase in cortical stiffness. To investigate the mechanisms underlying this stiffness increase, we probed individual cells after 30-minute treatment with different cytoskeleton-targeting drugs, namely 400 nM cytochalasin D (to depolymerize F-actin), 10 µM blebbistatin (to inhibit actomyosin contractility), and 10 µM Y-27632 (Rho-associated kinase (ROCK) inhibitor) (Figure 4C). All inhibitors caused a decrease in apparent Young’s modulus, consistent with the dominant role of actomyosin contractility in maintaining cortex tension.^[25]^ However, only F-actin depolymerization and inhibition of ROCK abolished the mechanical differences between cells originating from stiff and compliant hydrogels. Blebbistatin treatment significantly reduced stiffness of cells from both gels, but preserved the proportion of stiffness. ROCK is a major downstream effector of the small GTPase RhoA and plays important roles in cytoskeleton organization, e.g. by regulating F-actin dynamics.^[26,27]^ Taken together, our results following blebbistatin and Y-27632 treatment suggested that ROCK (independently from myosin) was responsible for the increased cell stiffness in stiff hydrogels, e.g. by altering the F-actin cytoskeleton. Consistently, we observed a marked increase in F-actin at the spheroids’ rim when grown in stiff hydrogels, while F-actin was more homogenously distributed in tumor spheroids growing in compliant hydrogels (Figure 4D). The cytoskeleton of luminal epithelial cells, like MCF-7 cells, consists predominantly of cytokeratin 8/18 (CK 8/18), known to be involved in maintaining cell integrity against mechanical stress.^[28]^ Cryo-sections revealed strong staining of the CK8/18 network throughout the spheroids (Figure 4D). Moreover, co-staining of F-actin and CK8/18 revealed higher levels of CK8/18 around the periphery of cells isolated from stiff hydrogels, and a slight decrease in staining intensity of the F-actin cortex (Figure 4E). Total levels of CK8/18 and F-actin were similar in Western blot and FACS analysis (Figure S7). Taken together, these results suggest that cytoskeletal reorganization was responsible for the seen changes in cell elasticity.

**Figure 4.**
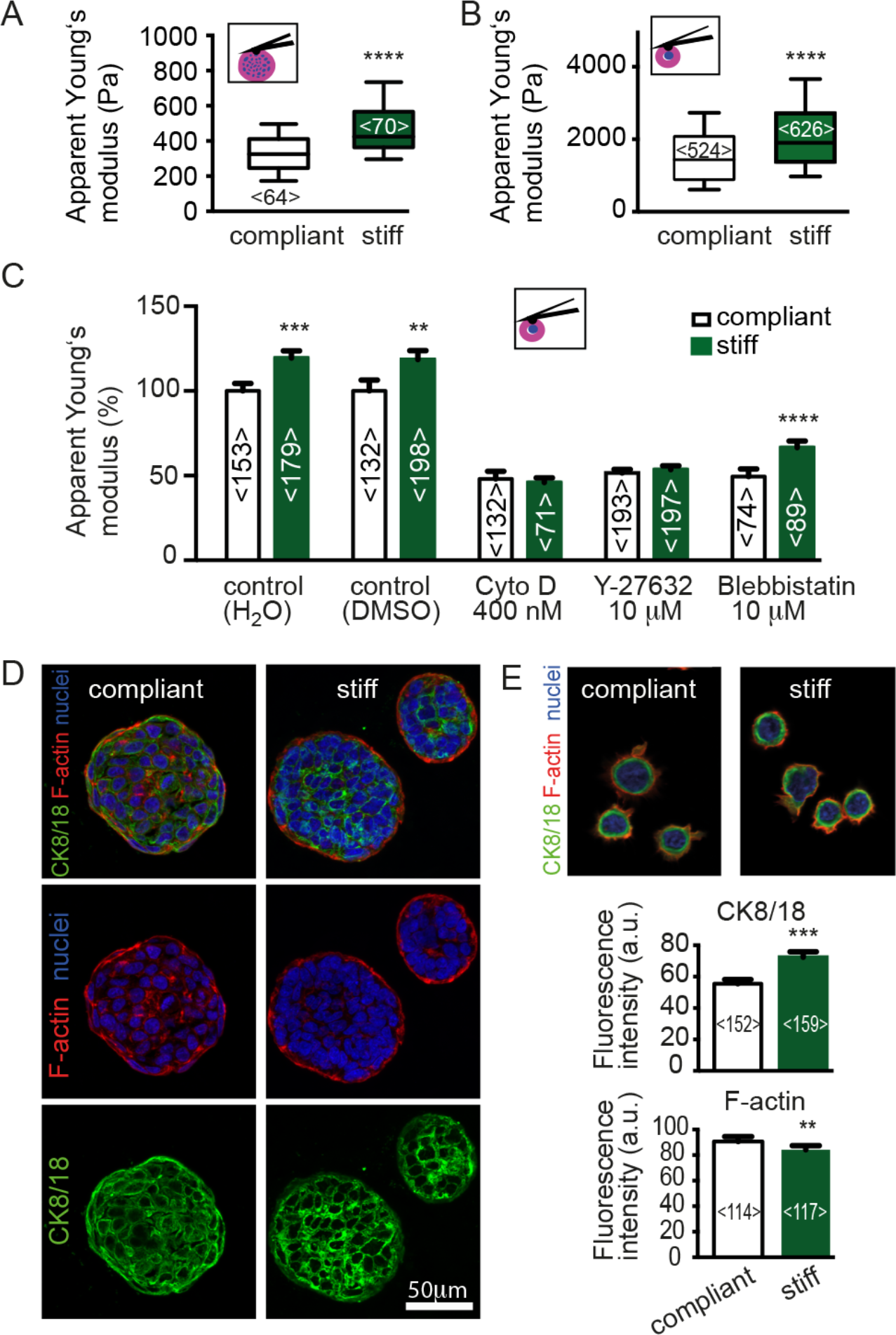
Characterizing the mechanical properties of tumor spheroids/isolated cells and their cytoskeleton. **A.** AFM indentation experiments on whole tumor spheroids. Tumor spheroids were released from compliant or stiff hydrogels and probed by AFM using a spherical indenter (diameter 10 µm). The number of probed spheroids is given **B.** AFM indentation experiments on single cells isolated from tumor spheroids and individually probed using a spherical indenter (diameter 5 µm). Apparent Young’s moduli in **A** and **B** are presented as Box-Whisker plots, where boxes mark the 25, 50 and 75 percentiles, and whiskers mark the 10 and 90 percentiles. **C.** Apparent Young’s moduli obtained by AFM indentation of individual cells isolated from compliant or stiff 3D cultures and treated for 30 min with drugs interfering with F-actin and actomyosin contractility before probing. Respective vehicle only controls (H_2_O for Y-27632) and DMSO (for cytochalasin D and blebbistatin) are shown. Data are normalized to respective controls (H_2_O for Y-27632 and DMSO for blebbistatin and Cytochalasin D. **D.** Immunofluorescence staining of cryosections of spheroids grown in compliant and stiff hydrogels. **E.** Immunofluorescence staining of single cells after isolation from 3D cultures (F-actin (red) and CK8/18 (green)). Bar graphs represent peak fluorescence intensities of radial profiles analyzed using image J (mean ± SEM). Numbers in brackets indicate the number of analyzed cells. Datasets compliant-stiff were compared using a Mann-Whitney test. **** denotes p-values < 0.001, *** p < 0.005, ** p < 0.01.

### Stiff microenvironments attenuate cell proliferation and increase p21 levels

Besides its role in cytoskeleton organization, ROCK has been implicated in regulating of proliferation of different cell types.^[29]^ Since we had observed differences in spheroid size (Figure 2B), we sought to investigate possible molecular mechanisms causing attenuated spheroid growth. Both, total metabolic activity (Figure 5A) and total DNA increase (Figure S8) were lower in stiff hydrogel cultures, which indicated diminished cell growth and was consistent with the reduced spheroid sizes (Figure 2). FDA/PI staining showed low numbers of dead cells, ruling out that decreased spheroid size was a result of increased cell death (Figure S9). This was confirmed by Western blot analysis of pro-apoptotic Bax, that showed no difference between stiff and compliant hydrogels (Figure S9). To investigate if hydrogel stiffness affected cell cycle progression, cell cycle distribution of Hoechst-stained cells was assessed by flow cytometry (Figure 5B). Stiff hydrogels elicited an increase in the percentage of cells in G0/G1 (on average 80% compared to 70% in compliant) and a decrease in the percentage of cells in S (10% versus 17%), G2 and M (9% vs 13% respectively). In line with this, we found a higher percentage of G1 (red) and a lower percentage of S/G2 (green) cells when we inspected MCF-7 FUCCI tumor spheroids (Figure 5C). These findings suggested that stiff hydrogels induced a G1 cell cycle delay or arrest.

**Figure 5.**
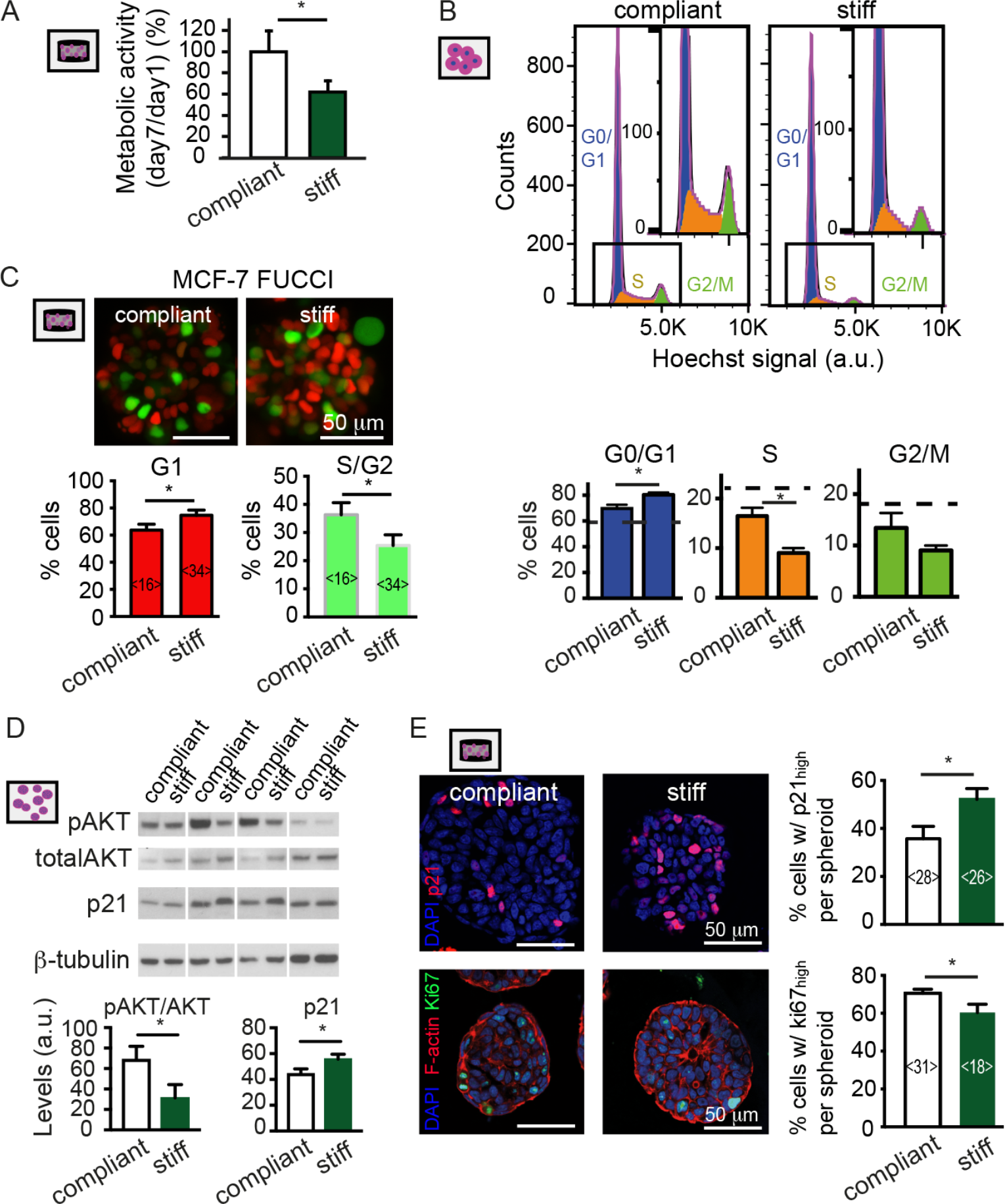
Analysis of cell cycle progression in dependence of hydrogel stiffness. **A.** Metabolic activity of tumor spheroid cultures was assessed using the Alamar Blue assay. **B.** Cell cycle analysis of Hoechst 33342-Stained cells isolated from tumor spheroids using flow cytometry. Histograms were fitted using the FlowJo Cell Cycle tool. Cell cycle stages are highlighted in different colors (3 independent experiments). **C.** Confocal images of MCF-7 FUCCI spheroids grown in compliant and stiff hydrogels. Bar graphs represent percentage of red (G1) and green (S/G2) cells (mean ± SEM). **D.** Western blot analysis of cell lysates from compliant and stiff hydrogel cultures for phospho-AKT (Ser473), total AKT and p21 (4 independent experiments). Bar graphs present densitometric quantification for pAKT/total AKT and p21 (mean ± SEM). p21 levels were normalized to ß-tubulin. **E.** Representative confocal images of immunofluorescently stained cryo-sections (above p21 (red)/nuclei (blue) and below Ki67 (green)/ F-actin (red)/ nuclei (blue)). Bar graphs represent nuclear staining intensities of p21 and Ki67, respectively, measured using FIJI. Distribution of intensity values was used to set a threshold defining high- and low-level fractions. The percentage of cells with high levels was determined per spheroid (mean ± SEM). Datasets compliant-stiff were compared using a t-test. * denotes p < 0.05.

Moreover, we investigated whether differences in the mechanical phenotype of compliant and stiff cultures could be linked to fluctuations in the mechanical properties exhibited along the cell cycle. We thus compared the mechanical properties of MCF-7 FUCCI cells by AFM and RT-fDC, a microfluidics-based technique for high-throughput cell mechanical characterization (Figure S15).^[30]^ Cells were more compliant in G1 than in G2 phase. Since cells in stiff cultures were stiffer and more frequently in G1 phase, we propose that cell stiffening is rather an effect of increased compression and not of cell cycle changes per se.

We next compared levels of cyclin-dependent kinase (CDK) inhibitors, p21 and p27, that play regulatory roles in the G1-S transition. Western blot analysis showed comparable levels of p27 (Figure S10), while p21 levels increased in spheroids cultured in stiff hydrogels (Figure 5D). Concomitantly, AKT phosphorylation at Ser^473^, an upstream regulator of p21, was reduced in spheroids grown in stiff hydrogels (Figure 5D). Nuclear p21 inhibits cell cycle progression via interaction with CDK2/4 and/or PCNA.^[31]^ Quantitative analysis of the nuclear staining intensities revealed an increased percentage of cells with high nuclear p21 levels in stiff hydrogel cultures (52% versus 36% in compliant hydrogels, Figure 5E), while the percentage of cells with high Ki67 levels was decreased in stiff hydrogel cultures (60% versus 71% in compliant hydrogels). Overall, these results show that microenvironment mechanics affected cell cycle progression.

### Modulating p21 and ROCK alters cell proliferation and mechanics

Since previous studies suggest a regulatory feedback loop between p21 and ROCK,^[31]^ we next set out to test if they were functionally linked in this context. We hypothesized that ROCK inhibition would affect not only cell stiffness (Figure 4C) but also tumor spheroid growth in 3D environments. To test this hypothesis, spheroids were grown in compliant and stiff hydrogels for 96 hours (starting from culture day 10) in the presence of ROCK inhibitor Y-27632 (or H_2_O controls). In both compliant and stiff hydrogels ROCK inhibition resulted in larger tumor spheroids (Figure 6A). Similar results were obtained for T47D tumor spheroids following addition of Y-27632 (Figure S11). Concomitantly, inhibition of ROCK in stiff hydrogels caused a reduction in the percentage of cells with high levels of nuclear p21, coupled with an increase in Ki67-positive cells (Figure 6B, C). Furthermore, single cells isolated from Y-27632 treated spheroids were significantly more compliant, although no inhibitor was added during mechanical characterization (Figure 6D). Consequently, ROCK inhibition reversed the stiff hydrogel phenotype and tumor spheroids resembled those cultured in compliant hydrogels, both in their mechanical properties and growth dynamics.

**Figure 6.**
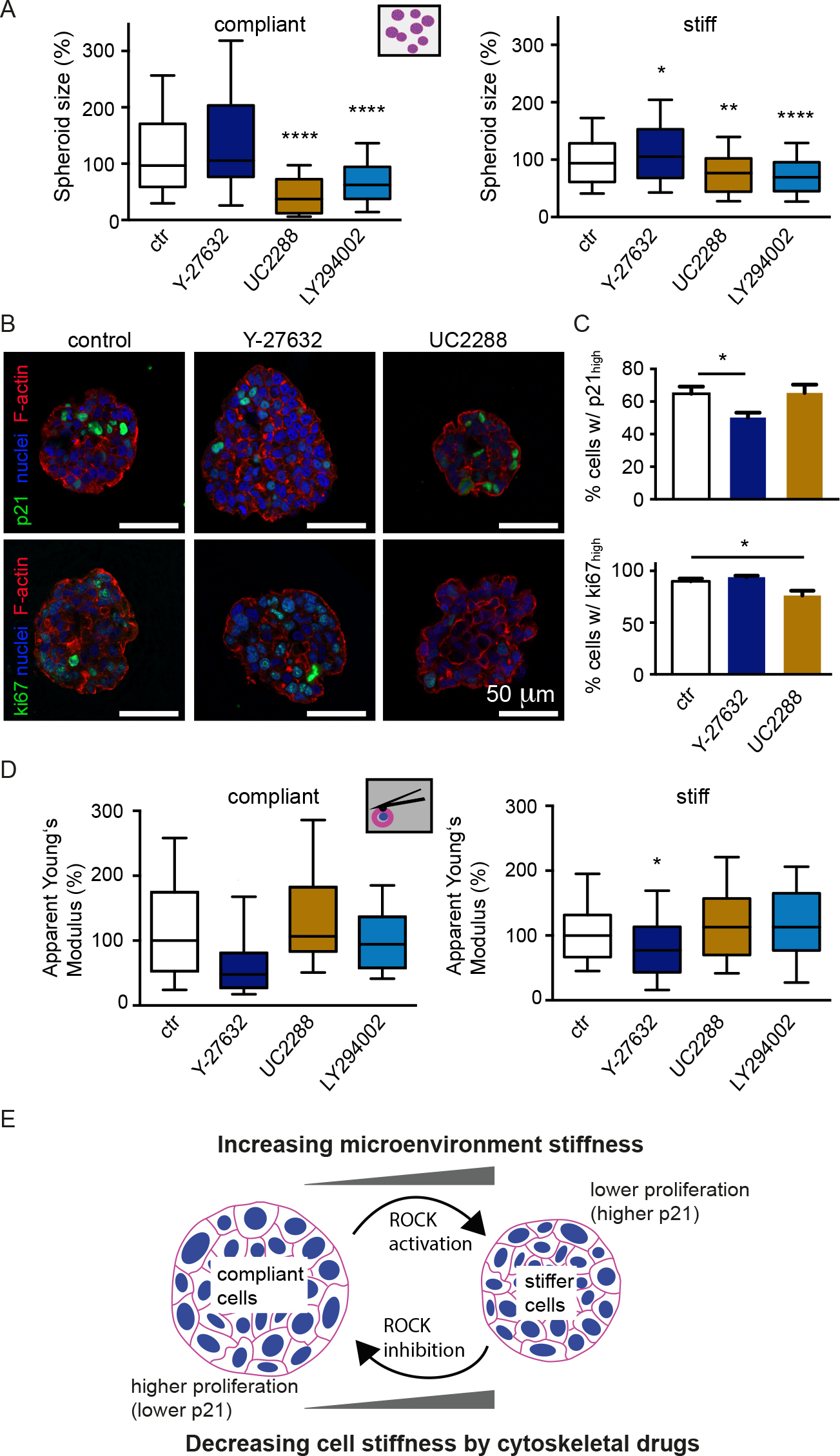
Testing the effect of ROCK and p21 inhibitors on spheroid growth and mechanics. **A.** Tumor spheroid sizes after 96-hour treatment with 10 µM Y-27632 (ROCK inhibitor), 2 µM UC2288 (p21 inhibitor) or 5 µM LY294002 (PI3K/AKT inhibitor). **B.** Confocal images of immunofluorescently stained MCF-7 cells grown in stiff hydrogels and treated for 96 hours with inhibitors **(A**). Cells were stained for total p21 or Ki67 (green), nuclei (blue) and F-actin (red). **C.** Nuclear staining intensities of p21 and Ki67, respectively, were measured in cryosections using FIJI. Distribution of intensity values was used to set a threshold defining high- and low-levels of expression. The percentage of cells with high levels was determined per spheroid. Datasets were compared using a one-way ANOVA with a Tukey’s multiple comparison test. * denotes p < 0.05. **D.** Bar graphs represent apparent Young’s moduli obtained by AFM indentation of individual cells isolated from compliant or stiff 3D cultures treated with respective inhibitors. Data in **A** and **D** are presented as Box-Whisker plots, where boxes mark the 25, 50 and 75 percentiles, and whiskers mark the 10 and 90 percentiles. Data in **A** and **D** were analyzed using a Kruskal-Wallis test. **** denotes p-values < 0.001, *** p < 0.005, ** p < 0.01, *p < 0.05. **E.** Summary illustrating the interrelationship between microenvironment stiffness, tumor spheroid growth and mechanics.

We next examined whether manipulating spheroid growth can affect the cell mechanical properties vice versa. The p21 inhibitor UC2288 has been previously described to reduce cytoplasmic p21 levels and to inhibit cancer cell growth.^[32]^ Both control and UC2288-treated cultures showed a dominant nuclear p21 staining with no detectable cytoplasmic p21 staining (Figure 6B, Figure S12). While we observed no significant difference in total and nuclear levels of p21 at 2 µM (Figure 6B), increasing concentrations of UC2288 (10 µM) resulted in elevated levels of nuclear p21 (Suppl. Figure 12). Since these higher concentrations of the inhibitor caused cell death (data not shown), we proceeded with a concentration of 2 µM. A 96-hour treatment with the inhibitor caused a significant decrease in spheroid size in both compliant and stiff gels (Figure 6A).

Meanwhile, we found no significant effect on the mechanical properties of cells isolated from spheroids (Figure 6C). Similarly, a 96-hour treatment with LY294002, a negative regulator of AKT activity, reduced spheroid growth as expected, but did not significantly affect the stiffness of released cells. Thus, we conclude that changes in the cell’s mechanical properties due to compression directly affected spheroid growth in a ROCK- and p21-dependent manner, while attenuated tumor spheroid growth itself did not affect cell stiffness (Figure 6E).

## Discussion

### Building a 3D model of growth-induced compressive stress

The mechanical properties of cancer cells and of their microenvironment are critical to breast cancer progression. How they are connected is not well understood. To address this question, we have investigated here the effect of microenvironment stiffness on the growth and mechanics of MCF-7 tumor spheroids. We made use of an engineered tumor microenvironment based on a synthetic PEG/heparin hydrogel, which allowed us to systematically tune microenvironment stiffness independently of ligand density. Additionally, we directly measured growth-induced stresses via the deformation of polyacrylamide beads/stress sensors embedded into the gel matrix. We quantitated compressive stresses of ~2 kPa at the spheroid edge in stiff degradable gels (Figure 3), which is comparable to previously reported stresses in tumor samples.^[10,33]^ For instance, several studies by the Jain lab have demonstrated that solid stress accumulates while tumors grow against the surrounding stroma.^[10,33]^ Some of this stress can be partly stored within the cell and matrix components of the tumor and is measurable after tumor dissection.^[33]^ For murine tumors, residual stresses in the range of 0.4-8.0 kPa, and for human tumors (including breast) stresses in the range of 2.2-9.0 kPa were determined.^[10,11]^ This approach, however, measures only residual stress stored in the tumor, while growth-induced stress balanced by the tumor microenvironment is not taken into account.^[33]^ Such growth-induced stresses were studied by the Jain lab using tumor spheroid cultures in non-degradable agarose gels. Theoretical growth-induced stresses were thereby calculated utilizing the spheroid size and mechanical properties of the gel.^[12]^ This theoretical approach relies on the assumption of a linear elastic gel deformation during spheroid expansion.^[12]^ When comparing the stresses calculated in our model to the range reported for the agarose gels (25-120 mmHg, ~6-16 kPa), lower values were found in our case (~2 kPa).^[12]^ Differences in the study design might account for the lower stress range observed here. In our model spheroids were cultured for 14 days, and their diameters did usually not exceed 150 µm, whereas Helmlinger et al. cultured spheroids for of up to 60 days, reaching diameters of up to 400 µm.^[12]^ Larger spheroids naturally cause higher stresses in the surrounding matrix. Another important difference between the models is, that our PEG/heparin hydrogels feature matrix degradation sites that enable cells to locally degrade the hydrogel during tumor spheroid growth, thereby creating space for the expanding tumor spheroid volume. Consistently, we observed that spheroids reached at comparable stresses larger sizes in degradable compared to non-degradable gels (Figure 3). Thus, growth reduction due to compression may start in non-degradable gels at much smaller spheroid sizes. Interestingly, no bead compression was evident in compliant degradable (unlike compliant non-degradable) hydrogels, suggesting that the compliant hydrogel matrix was degraded faster than the spheroid grew (Figure 3). It was recently reported for fibroblasts that MMP release correlates inversely with matrix stiffness,^[34]^ which may suggest that in degradable compliant hydrogels an increase in MMP release additionally contributed to stress relieve. Since matrix degradability is an important aspect of the real tumor microenvironment, polyacrylamide stress sensor beads embedded in the hydrogel matrix are a straightforward method to monitor growth-induced stresses in the tumor spheroid environment. Several groups have recently used polymeric beads as stress sensors to measure stresses acting within tumor spheroids. For example, polyacrylamide beads were embedded as local stress sensors within multicellular aggregates.^[35]^ Another study used alginate beads loaded with fluorescent nanoparticles to measure compressive stresses within melanoma cell spheroids.^[36]^ To our knowledge, polymeric stress sensors have not been used before to measure growth-induced stress in the environment of a tumor spheroid. The novel approach taken in our study therefore provides valuable insights into how tumor spheroids balance their growth and matrix degradation in response to environment stiffness.

### The effect of stiff hydrogel cultures on spheroid growth

In our study, we observed that stiff microenvironments negatively influence tumor spheroid growth (Figure 2). Such effect of compressive stress on spheroid size has been previously reported for tumor spheroids assembled from human colon, murine mammary carcinoma, and rat sarcoma cells growing in agarose gels,^[12]^ and breast cancer cells (MCF-7) in alginate gels.^[37]^ This was attributed to an increased cellular packing density, but not to a decrease in proliferation nor increase in apoptosis.^[12]^ In fact, decreased levels of apoptosis were detected in stiff hydrogels, which was interpreted as result of the increased cell-cell interactions in more compact spheroids.^[12]^ In accordance to these findings, we also detect an increased packing density of cells and no increased levels of apoptosis in tumor spheroids growing in stiff hydrogels (Figure 2G, Figure S9). Conversely, we found decreased proliferation in stiff hydrogels, which was associated with delay/arrest of cell cycle progression (Figure 5). Thus, we attribute the reduced tumor spheroid sizes to both increased compaction and lower proliferation.

To investigate the reasons accompanying increased packing density, we used TEM to look for voids or gaps between cells. We observed no intercellular spaces but occasionally pores throughout the gels that may collapse upon compression and contribute to increased cell density. Alternatively, cell volume reduction may cause volume reduction of tumor spheroids. While after isolating the cells from the spheroids no significant difference in average cell size was seen (Figure S13), cell volumes might have been reduced due to their compression and water efflux in intact spheroids. This would be consistent with the slightly decreased nuclear sizes (9%, Supplementary Figure 2B). Osmotically induced volume changes occur at much shorter timescales than it takes to release spheroids from the hydrogels (seconds or even less) ^[38]^, which would explain why eventual volume changes were not detected after isolation of single cells. Cell volume changes due to water efflux can occur under isotonic conditions, as observed during cell spreading on stiff 2D substrates, which requires increased ion channel activity to export osmolytes.^[39]^

In accordance with our findings, several studies reported a G1 cell cycle arrest/delay upon compression. For instance, in microbes under confinement,^[40]^ 2D cultures of MDCK cells growing on stretchable membranes,^[41]^ and in sarcoma, colon and breast spheroids that were osmotically compressed by dextran addition to their cell culture medium.^[13]^ The latter study suggested that osmotic compression inhibits spheroid growth via p27,^[13]^ another CDK inhibitor of the same family as p21. Analogously, we found increased levels of nuclear p21 in spheroids isolated from stiff hydrogels. A dual role for p21 in cancer progression has been reported before.^[42]^ Phosphorylation of p21 by AKT attenuates its interaction with CDK 2/4 and PCNA and thereby promotes cell cycle progression.^[43,44]^ In line with this, we observed lower levels of phospho-AKT concomitant with increased nuclear p21 and cell-cycle arrest/delay in G1. Thus, our data suggest that increased levels of nuclear p21 attenuate spheroid growth in stiff hydrogels.

### The effect of stiff hydrogel culture on cancer cell mechanics

Stiffness values of both whole spheroids and individual cells were significantly higher when spheroids were cultured in stiff hydrogels. This increased cell stiffness reflects changes in cytoskeletal organization, which were shown to be dependent on F-actin and ROCK (Figure 4). Increased ROCK activity due to compression would be in line with a recent study on 3D cultures and tissue explants, which demonstrated that acute compression directly activates the Rho/ROCK pathway.^[45]^ While several studies reported that elevated substrate stiffness can increase the stiffness of various cell types, including human mesenchymal stromal cells in 2D cultures,^[46,47]^ little is known about the effect of the mechanical properties of a 3D environment on cell mechanics. Byfield et al. have found that the stiffness of endothelial cells increases in stiff collagen gels.^[48]^ The Zaman lab compared the mechanical properties of breast and prostate cancer cells upon culture in 3D collagen gels of varying stiffness by microrheology and reported a respective increase or decrease in cell stiffness in stiffer gels.^[49][50]^ Malignant cancer cells were also recently found to adapt their mechanical properties to their surrounding.^[51]^ In the aforementioned studies, however, only single cells and no tumor spheroids were investigated, so the effect of growth-induced compressive stress on cell mechanics was, to our knowledge, not addressed before. We also investigated cytoskeletal candidates that may be responsible for the increased cell stiffness. Accumulation of F-actin was observed at the rim of tumor spheroids grown in stiff hydrogels. Additionally, cells isolated from stiff hydrogels displayed increased CK8/18 but decreased F-actin staining intensities. The increase in CK8/18 might be associated with its reported role in protecting tissues and cells against mechanical stress.^[28]^ Decreased F-actin staining levels in individual cells isolated from stiff hydrogels correlated with increased stiffness values, which appears at first counter-intuitive. Since we indented cells typically less than 0.5 µm in our AFM measurements, we expect to probe predominantly the mechanical properties of the F-actin cortex. However, differences in staining intensities do not necessarily reflect the mechanical properties of the cortex, since the network stability is largely dependent on crosslinking, other associated proteins, and their spatial organization.^[52]^ Overall, changes in cell mechanics and relative levels of cell cytoskeleton components suggest that cells reorganized their cytoskeleton according to compressive stress.

### The role of ROCK in linking cell proliferation and mechanics

ROCK activity attenuated spheroid growth via p21, as shown by the ROCK dependent increase in cell elasticity in stiff hydrogels, increased growth of spheroids in presence of ROCK inhibitor, and decreased levels of p21. It may be possible that the observed cytoskeletal changes, seen in our system, reflect a mechanism regulating cell proliferation under compression, which breaks down in case of ROCK inhibition. Various studies proposed a functional link between ROCK, AKT, and p21. ROCK inhibition can promote AKT phosphorylation,^[53]^ and AKT is a well-known regulator of p21.^[31,43]^ Thus, in our stiff hydrogels increase in ROCK activity (Figure 4C) may have attenuated spheroid growth via lower AKT phosphorylation and p21 regulation (Figure 5). Moreover, a feedback loop has been suggested, where p21 itself can influence ROCK activity.^[31]^ To test this possibility, we used an inhibitor of p21, UC2288, that has been previously described to reduce total and cytoplasmic but not nuclear p21 levels.^[32,54,32]^ Treatment of tumor spheroids with moderate concentrations of UC2288 did not significantly affect total p21 levels, but decreased cell proliferation as indicated by lower Ki67 levels and reduced spheroid sizes in both stiff and compliant gels (Figure 6C). This is in line with UC2288’s reported role in attenuating growth of different cancer cells.^[31,32]^ Conversely, UC2288 had no effect on the mechanical properties of cells. From this we conclude that in our system ROCK activation due to compression caused cellular stiffness increase and elevated nuclear p21 levels but was not a result of p21 modulation.

ROCK inhibition resulted in larger spheroids that were, however, less compact compared to controls (Figure S14). This was unexpected since larger tumor spheroids should experience increased levels of compressive stress. Since we detected lower cell stiffness after Y-27632 treatment (Figure 6D), it appears that cells within tumor spheroids can handle compressive loads, at least to a certain extent, without reinforcing their F-actin cortex. A possible mechanism may consist in osmotic regulation, as previously reported for MCF-7 spheroids growing in agarose gels.^[55]^ Osmotic regulation is associated with F-actin cytoskeletal reorganization and involves the RhoA/ROCK pathway as previously described.^[56]^ Besides osmotic stress, it is well known that the Rho/ROCK pathway can be activated by different extracellular stimuli, and its effect on cell proliferation is highly cell type- and context-dependent.^[29]^ While the majority of studies report that interfering with ROCK (e.g. by Y-27632) decreases proliferation, ^[57]^ others found increased proliferation, ^[53,58]^ in line with our results. For instance, Nakashima et al. have described a mechanism by which ROCK negatively regulates EGF-induced colon cancer cell proliferation.^[58]^ Increased expression and activity of ROCK has been associated in various studies with cancer progression.^[29]^ For example, higher ROCK activity in MCF-7 cells increased their metastatic potential in a mouse model of breast cancer bone metastasis. ^[59]^ There are, however, also studies that attribute ROCK a role as negative regulator of cancer progression.^[29]^ Thus, given these controversial effects of ROCK inhibition, clearly a more in-depth investigation in different cellular and environmental contexts is needed before considering ROCK as potential therapeutic target.

### Cell mechanics and cancer progression

It is well accepted that the cell’s mechanical properties are altered during cancer progression.^[60,61]^ To date, more than 25 studies have compared cell stiffness of different types of cancer cells (breast, prostate, kidney, colon etc.) of varying invasiveness and mostly agree that more invasive cancer cells are more compliant. This may be an important aspect during cancer cell invasion, where disseminated tumor cells migrate in confined spaces of the tumor stroma.^[62]^ At this stage, however, it is not clear if invasive cancer cells are also more compliant in a tissue context. We have found here that MCF-7 cells adapt their mechanical phenotype upon tumor spheroid culture in hydrogels of different stiffness. Although in compliant hydrogels single cells were more compliant compared to stiff hydrogels, they did not show any clear signs of increased invasiveness. It is possible that the effect of microenvironment stiffness on tumor cell mechanics is cell-type dependent and different factors, such as increased ligand density or additional geometric cues (e.g. fibrillar arrangements) are required. Nevertheless, we show here that stiffer microenvironments caused increased compressive stress, increased cell stiffness and reduced tumor spheroid growth, which are all features known to counteract tumor progression.

## Conclusions

To summarize, we describe here a mechanism involving ROCK and p21, by which compression, due to growth under confinement in stiff environments, regulates tumor spheroid growth by altering the cytoskeletal/mechanical status of constituent cells. Together, this may present a regulatory mechanism, by which cells decrease their proliferation, when they encounter increasing compression. Nonetheless, the upstream mechanism by which compression is initially sensed to modulate the RhoA/ROCK pathway is not yet identified and requires further exploration. Since both the mechanical properties of cancer cells and also of their microenvironment are critical to cancer progression, a better understanding of their link may help to develop new therapeutic approaches for the treatment of cancer.

## Methods

### Cell culture

MCF-7 cells were obtained from the “Deutsche Stammsammlung für Mikroorganismen und Zellkulturen” (DSMZ) and maintained according to their recommendations in RPMI 1640 (Roswell Park Memorial Institute) supplemented with 10% heat-inactivated fetal bovine serum (FBS), 1 mM sodium pyruvate, MEM non-essential amino acids, 10 µg/ml human insulin (Sigma) and penicillin (100 U/ml) -streptomycin (100 µg/ml). MCF-7 FUCCI cells, recently described and characterized ^[63]^, were kindly provided by A. Ferraris lab. MCF-7 FUCCI cells were cultured in DMEM (Dulbecco’s Modified Eagle Medium) supplemented with 10% heat-inactivated fetal bovine serum (FBS) and penicillin (100 U/ml) -streptomycin (100 µg/ml). Before starting 3D cultures, cells were grown in standard T75 culture flasks and sub-passaged 2-3 per week. For cell detachment, TrypLE^®^ was used. All cell culture reagents were from Thermofisher unless otherwise stated.

### Preparation of PEG-heparin 3D hydrogel cultures

PEG and heparin-maleimide precursors were synthesized as previously described.^[22,23]^ Cells were detached from tissue culture flasks with TrypLE^®^, resuspended in cell culture medium, centrifuged down, and resuspended in PBS. Then they were mixed with the freshly prepared heparin-maleimide (MW~15000) solution at a final density of 5.6 × 10^5^ cells/ml and a final heparin-maleimide concentration of 1.5 mM. For a γ of 1.5 (molar ratio PEG/heparin-maleimide), PEG precursors (MW~16000) were reconstituted with PBS at a concentration of 2.25 mM and placed into an ultrasonic bath (Merck) for 20 s (medium intensity). Lower stiffness gels were prepared by diluting the stock solution of PEG precursors accordingly (1.5 mM for γ=1.0, 0.75 mM for γ=0.75). To prepare PEG-heparin hydrogel droplets, heparin-cell suspension was mixed in a chilled microtube with the same amount of PEG-solution using a low binding pipette tip. Then, a 25 µl drop of the PEG-heparin-cell mix was pipetted onto hydrophobic, Sigmacote^®^ (Sigma Aldrich)-treated glass slides (VWR). Gel polymerization started immediately and stable hydrogels were obtained within 1-2 minutes. Hydrogels were gently detached from the glass surface after 4 minutes using a razor blade and transferred into a 24 multi-well plate supplemented with 1 ml cell culture medium. Cell culture medium was exchanged every other day. For experiments with inhibitors UC2288 (2, 4, and 10 µM) (Merck), LY294002 (5 µM) (Sigma Aldrich) and Y-27632 (10 µM) (Sigma Aldrich), inhibitors (or DMSO/H2O controls respectively) were added on day 10 for a 96-hour treatment.

### Quantification of hydrogel mechanical properties by AFM

After preparation, hydrogels were immersed in PBS and stored at 4 °C until mechanical characterization on the following day. After equilibrating them at RT for 1 hour, gels were mounted using CellTak (Thermofisher) onto glass object slides (VWR). A Nanowizard I or IV (JPK Instruments) was used to probe the gel stiffness. Cantilevers (arrow T1, Nanoworld), that had been modified with polystyrene beads of 10 µm diameter (Microparticles GmbH) using epoxy glue (Araldite), were calibrated using the thermal noise method implemented in the AFM software. Hydrogels were probed at RT in PBS using a speed of 5 µm/sec and a relative force setpoint ranging from 2.5 to 4.0 nN in order to obtain comparable indentation depths. Force distance curves were processed using the JPK data processing software. Indentation parts of the force curves (approximately 1.5 µm indentation depth) were fitted using the Hertz/Sneddon model for a spherical indenter, assuming a Poisson ratio of 0.5.^[64,65]^

### Isolation of spheroids and single cells from hydrogel cultures

On day 12-14 of culture, spheroids were harvested by gel degradation. Briefly, hydrogels were washed once with PBS (with Mg^2+^, Ca^2+^). Then, gels were incubated with 1 ml 2.5 mg/ml collagenase (NB 4G, Serva) for 16-18 min at 37 °C. Then, 5 ml PBS was added and gels/spheroids were centrifuged (80g, 3 min). These steps resulted in a pellet of isolated and intact spheroids that were then either directly lysed for Western blot analysis, used for AFM characterization, or processed for further isolation of single cells. To further degrade the spheroids to obtain a single cell suspension, spheroids were incubated with TrypLE for 10 min at 37 °C. Cells were then washed in cell culture medium, spun down at 160g for 4 min and resuspended in CO_2_-independent medium.

### Morphometric analysis of tumor spheroids *in situ* and *ex situ*

At day 14, cultures were fixed with 4% formaldehyde/PBS for 1 hour, followed by a 30 min permeabilization step in 0.2% Triton-X100. Spheroids were stained for 4 hours with 5 µg/ml DAPI and 0.2 µg/ml Phalloidin-TRITC in 2% BSA/PBS. Then hydrogels were immersed at least for 1 hour in PBS. For imaging, hydrogels were cut into slices with a blade and were placed on a cover slide to image the interior of the hydrogel. Spheroids were imaged with a LSM700 confocal microscope using a 40x objective (Zeiss C-Apochromat). Quantitative analysis of cell morphology was done using FIJI. Briefly, a threshold was applied to the fluorescent images (red channel, F-actin), which were then transformed into binary pictures, and shape factor (defined as 4 × π × area/(perimeter)^2^) and area were calculated. To analyze the diameters of a larger number of tumor spheroids, they were isolated from the hydrogels as described above and imaged using phase contrast microscopy. The diameters were then measured manually using FIJI. For analysis of MC7-7 FUCCI spheroids, z-stacks of spheroids (77 slices at steps of 1 µm) were recorded with a LSM700 confocal microscope using a 20x objective. From maximum intensity projections the number of red/green cells was determined. At least 16 spheroids were analyzed per condition.

### AFM characterization of single cells or spheroids

A Nanowizard 1/4 (JPK Instruments) was used to probe the stiffness (apparent Young’s modulus) of entire spheroids or cells isolated from these. Cantilevers (arrow T1, Nanoworld) modified with polystyrene beads with a diameter of 5 µm (for single cells) or 10 µm (for whole spheroids), were calibrated using the thermal noise method implemented in the SPM software. Spheroids or single cells suspended in CO_2_-independent medium were pipetted into glass bottom dishes (World Precision Instruments). Once the cells/spheroids had settled and were stably attached to the surface of the dish, the cantilever was lowered onto the top of each spheroid/cell at 5 µm/sec speed. Isolated cells were probed in their center using a relative force set-point of 2.5 nN. At least 50 cells per condition were probed in each experiment. In the case of spheroids, a grid of 20 µm × 20 µm (4 × 4 points) was set and 10 - 20 spheroids were probed in each experiment using a relative force set-point of 5 nN. Force distance curves were processed using the JPK data processing software for indentation depths of 10-20% using the Hertz/Sneddon model for a spherical indenter and assuming a Poisson ratio of 0.5.^[64–66]^ For single cells, a correction factor accounting for the bottom deformation of the cell was taken into account as suggested previously. ^[67]^

### Preparation of polyacrylamide hydrogel beads

Polyacrylamide (PAA) hydrogel beads were produced as recently described using a microfluidic droplet generator.^[24]^ An average Youngs modulus of approximately 4 kPa was determined using AFM. To visualize beads within the hydrogels, beads were functionalized with Poly-L-Lysine (PLL) conjugated with Cy3 fluorophores as described before.

### Quantification of bead deformation

From confocal z-stack orthogonal views, the 3D distance between a particular bead and the spheroid edge was determined in the Zen Software (Zeiss). The aspect ratio of the beads was measured using Fiji in the maximum projection. Only beads located within a narrow angular range with respect to the equatorial plane (± 20°) were used for analysis, since otherwise bead deformation was underestimated in maximum projections. From the aspect ratio, the resultant radial stress component was calculated as described below. Aspect ratio and radial pressure were plotted versus the distance (bead-spheroid edge) using Igor Pro (Wavemetrics).

### Calculation of compressive stress acting on tumor spheroids

Consider a tumour spheroid that generates a rotationally symmetric displacement field in a surrounding incompressible hydrogel. Because of incompressibility, we will only consider the traceless part of the stress tensor.

Due to rotational symmetry, we have for the traceless components of the stress tensor^[68]^

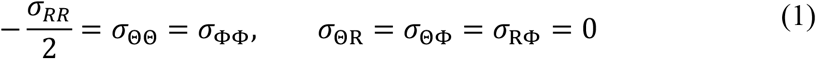

where *R* and Φ are spherical coordinates of the coordinate system centred at the spheroid midpoint.

Consider now an incompressible bead at position <u>*R*</u>_*bead*_ in the neighbourhood of the spheroid that is being deformed due to the stress field in the gel surrounding the tumour spheroid. In the following, we will relate the aspect ratio of a slightly deformed spherical bead to the radial stress component *σ*_*RR*_ using linear elasticity theory. As the bead radius *r*_*bead*_ is considerably smaller than the spheroid radius, we can make the approximative assumption that the bead feels a stress field which is axisymmetric where the symmetry axis is the line connecting the spheroid and the bead centre. Furthermore, we will assume that the stress field varies to a negligible degree along the length scale of the bead diameter. Let (*φ*, *z*, *ρ*) be cylinder coordinates of the reference frame with origin at the bead center and the z-axis coinciding with the radial direction set by the tumor spheroid. Then, *σ*_*zz*_ ≈ *const*. ≈ *σ*_*RR*_(<u>*R*</u>_*bead*_). Due to Eq. (1), we have

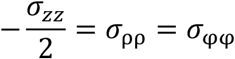

Let (*r*, *θ*, *φ*) be spherical coordinates with respect to the bead reference frame.

We have

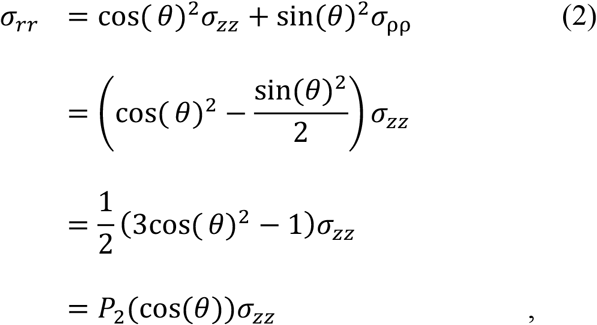

where *P*_2_ is the Legendre polynomial of 2nd order.

Furthermore, we have

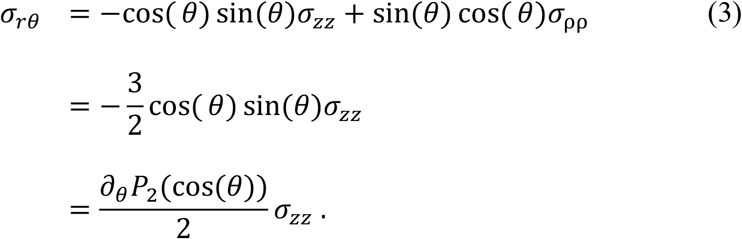

For an axisymmetric problem, a general solution that solves the force balance condition

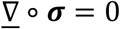

is given by the displacement field ^[69,70]^

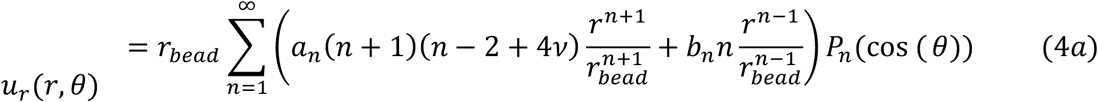

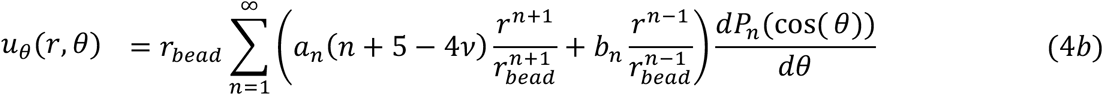

with associated stress components

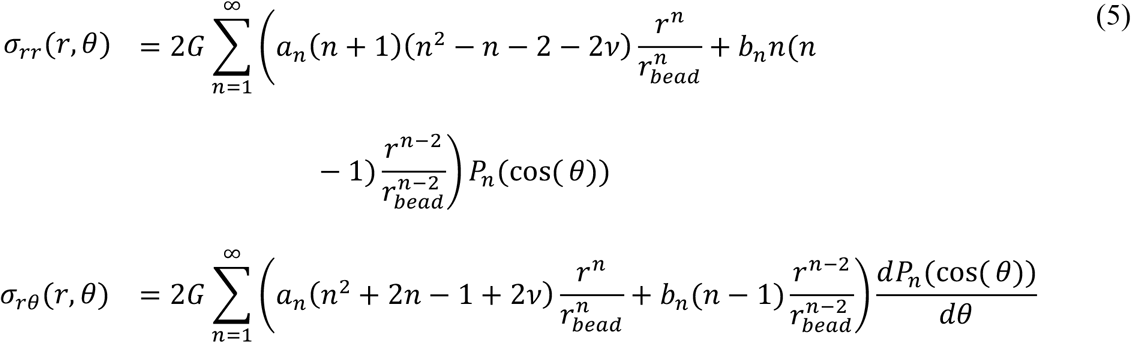

From Eqn. (2) and (3), we conclude that only contributions for n=2 are relevant in Eqn. 5 with *a*_2_ = 0 and *b*_2_ = *σ*_*zz*_/(4*G*). For the deformation field on the bead surface, we get with Eq. 4a

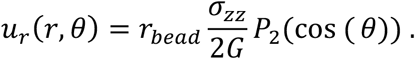

To first order, the aspect ratio *AR* of the bead is given by

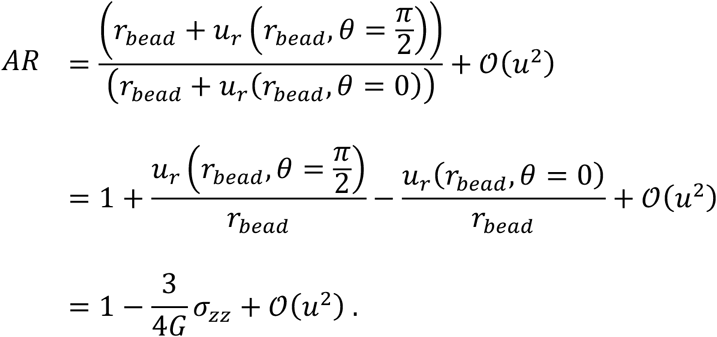

Therefore,

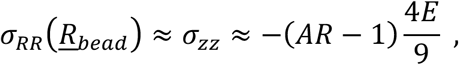

where we used that *G* = *E*/3 for the incompressible case. Note that for the case of a growing tumor spheroid, the deformed gel exerts a positive pressure on the spheroid giving rise to a negative value of *σ*_*RR*_ and an oblate shape of the bead with aspect ratio AR>1.

### Western blot analysis

For Western blot analysis, spheroids were isolated from hydrogels as described above and lysed using Laemmli sample buffer (62.5 mM Tris-HCl (pH 6.8), 2% SDS, 10% glycerol, 0.01% bromophenol blue). For analysis of cytokeratins, a different cell lysis protocol was used to obtain Triton X-100-soluble and insoluble fractions as described elsewhere.^[71]^ Samples were boiled (5 min 95 °C), loaded on gradient gels (MiniProtean, Biorad), and separated by reducing SDS-PAGE. After blotting onto a PVDF membrane (Merck Millipore), membranes were blocked with TBS-Tween (20 mM Tris, 137 mM NaCl, 0.1% Tween) containing 5% (w/v) non-fat dry milk for 1 hour. Membranes were incubated overnight at 4°C (5% BSA/PBS) with antibodies against total p21 (12D1, Cell Signaling), tubulin (Abcam), GAPDH (Abcam), pAKT (Ser473, Cell Signalling), panAKT (Cell signalling), p27 kip1 (Cell Signaling), cyclin D1 (Dianova), CK8/18 (Dianova, Abcam). After incubation with respective HRP-conjugated secondary antibodies (Abcam), chemiluminescence detection was performed using Enhanced Chemi-Luminescence (ECL) (Thermofisher). Densitometric analysis was done using Fiji.

### Immunofluorescence staining of cryosections

After fixation, hydrogels were transferred into 30% of sucrose/PBS for 1 hour, and thereafter embedded into optimum cutting temperature cryo-embedding compound (O.C.T., Thermofisher) and snap frozen in liquid nitrogen. 3D samples were sectioned in 8 – 10 µm thick cryosections with the Microm HM560 Cryostat (Thermofisher). Antigen retrieval was performed by immersing slides into citrate buffer (0.1 M, pH 6.0)/ 0.05% Triton X-100 (Sigma) for 10 min. Then slides were incubated for 10 min in 0.1% Triton X-100/PBS and blocked for 1 hour in 2.5% BSA/PBS. Slides were incubated with primary antibodies in 2.5% BSA/PBS overnight at 4 - 6°C in a humidified chamber. Following primary antibodies were used: p21 (Cellsignaling), CK8/18 (Dianova), and ki67 (Dianova). After washing the glass slides with PBS, slides were incubated for 2 hours with secondary antibodies (Cy5-conjugated anti-mouse/rabbit IgG, Dianova) and Phalloidin-TRITC (Sigma) and DAPI (Sigma). Finally, the slides were washed with PBS and mounted after a short wash in ddH_2_O with mounting medium (Thermofisher). Samples were imaged with a LSM700 confocal microscope (Zeiss) using a 40x objective (C-Apochromat, Zeiss). For quantification of nuclear staining intensity levels of p21 and ki67, the DAPI channel was selected and individual nuclei were segmented: first, using the “find maxima-segmentation” function in Fiji, individual nuclei were segmented. A duplicate of the DAPI image was generated and after applying a threshold, it was transformed into a binary image. Both images were then recombined using the “image calculator- AND” function. The segmented image was used to define regions of interest. Redirecting to the channel showing the p21 or ki67 staining (Cy5) using the “Set Measurements” function, their respective staining intensity within the nuclear region was calculated using the “Analyse Particles” function.

### Measurements of metabolic activity using Alamar Blue^®^ assay

At day 0, 4 and 7, cell culture medium was removed from the hydrogels and 300 µl of 6% Alamar Blue^®^ (Thermofisher) solution in serum-free medium were added. After incubation for 3 hours at 37 °C, 100 µl of the supernatant were transferred in triplicate into a black 96 well plate. Fluorescence emission was measured at 590 nm on a plate reader (Tecan). Data for day 4 and day 7 were normalized against day 0. Three gels of each condition were analyzed in each experiment and the experiment was repeated twice.

### Flow cytometry on cells isolated from tumor spheroids

Cells were isolated from spheroids as described above, fixed for 5 min with 4% formaldyhyde/PBS, permeabilized with 0.2% Triton X, and stained with antibodies against CK8/18 (Dianova). To analyze cell cycle distribution of cells, cells were stained with 8 µM Hoechst 33342. Cells were analyzed using a flow cytometer (LSR II, Becton Dickinson). Cell cycle profiles were analyzed using the “Cell cycle analysis” tool in FlowJo (FlowJo).

## Statistical analysis

GraphPad Prism was used to plot data and to perform statistical tests. A two-tailed significance level of 5% was considered statistically significant (p<0.05). The respective tests applied for differences between independent groups (e.g. compliant versus stiff), are indicated in figure legends. In principle Mann-Whitney (non-parametric) or t-test (in case of normal distribution) were used to compare two independent datasets. In case of more than two groups, a Kruskal-Wallis (non-parametric) or one-way ANOVA (normal distribution) test with multiple comparisons (Tukey) were chosen. To test for normality, a D’Agostino-Pearson omnibus normality test was performed. Statistical analyses were carried out using 1D linear mixed model that incorporates fixed effect parameters and random effects to analyze differences between cell subsets and replicate variances, respectively. *p*-values were determined by a likelihood ratio test, comparing the full model with a model lacking the fixed effect term.

## Acknowledgements

We thank all Guck lab members, in particular members of the cell and tissue mechanics group and Dr. Angela Jacobi, Dr. Philipp Rosendahl and Dr. Oliver Otto for scientific discussions and help with experiments. Many thanks to Dr. Despina Soteriou for revising the manuscript draft. We thank JPK Instruments for technical support. We are also thankful to Prof. A. Ferrari’s group for providing MCF-7 FUCCI cells. This project has received funding from the Alexander-von-Humboldt Stiftung (Humboldt-Professorship to J.G.). We thank the microstructure, electron microscopy, and flow cytometry facility of the CMCB (in part funded by the State of Saxony and the European Fund for Regional Development-EFRE).

## Abbreviations

2D/3D: two/threedimensional
AFM: atomic force microscopy
ECM: extracellular matrix
CDK: cyclin dependent kinase
CK8/18: cytokeratin 8/18
MCF-7: Michigan Cancer Foundation-7
PEG: polyethylene glycol
MMP: matrix metalloproteinase
TEM: transmission electron microscopy
SEM: scanning electron microscopy
PBS: phosphate buffered saline

