## Supplemental Information for "3D microenvironment stiffness regulates tumor spheroid growth and mechanics via p21 and ROCK"

### Supplementary Methods

#### Transmission Electron Microscopy (TEM)

Tumor spheroids within hydrogels were fixed in modified Karnovsky's fixative (2% glutaraldehyde + 2% paraformaldehyde in 50 mM HEPES) for at least overnight at 4°C (1). Samples were washed twice in 100 mM HEPES and 2x in water and postfixed/contrasted by treatment with osmium tetroxide ($OsO_4$), thiocarbohydrazide (TCH), and again $OsO_4$ (O-T-O, (2, 3)). In brief, samples were incubated in 2% aqueous $OsO_4$ solution containing 1.5% potassium ferrocyanide and 2 mM $CaCl_2$ for 30 min on ice. After washes in water, the samples were incubated in 1% TCH in water (20 min at room temperature), followed by washes in water and a second osmium contrasting step in 2% $OsO_4$/water (30 min, on ice). After that, samples were washed in water, en bloc contrasted with 1% uranyl acetate/water for 2 hours on ice, washed again in water, dehydrated in a graded series of ethanol/water (30%, 50%, 70%, 90%, 96%, 3x 100% ethanol (pure ethanol on molecular sieve)), and infiltrated in epon 812 (epon/ethanol mixtures: 1:3, 1:1, 3:1 for 1.5 hours each, pure epon overnight, pure epon 5 hours). Finally, the samples were embedded in flat embedding molds and cured at 65°C overnight. Ultrathin sections were prepared with a Leica UC6 ultramicrotome (Leica Microsystems, Vienna, Austria), collected on formvar-coated slot grids, and stained with lead citrate and uranyl acetate. Contrasted ultrathin sections were analysed on a FEI Morgagni D268 (FEI, camera: MegaView III, Olympus) or a Jeol JEM1400 Plus (JEOL, camera: Ruby, JEOL) both at 80 kV acceleration voltage.

**Assessing cell growth using PicoGreen® assay**

Hydrogels were transferred into 1.5 ml microtubes and frozen at −80 °C. After thawing, samples were incubated overnight at 37 °C with 300 µl of 10 U/ml Proteinase K (Sigma Aldrich) in PBS-EDTA, followed by 8 hours at 56°C. Samples were then treated four times for 10 min in a 4°C cold ultrasonic bath. DNA was quantified using the PicoGreen® assay according to the manufacturers' instructions. Values for day 4 and day 7 were normalized to day 0.

**Real-time fluorescence and deformability cytometry (RT-FDC)**

RT-FDC on MCF-7 FUCCI cells was performed as described previously (4). In brief, MCF-7 FUCCI cells (5) were detached and resuspended in PBS containing 0.6% methylcellulose to a final concentration of approximately $3x10^6$ $mL^{-1}$. The cell suspension was drawn into a syringe and flushed through a 30 µm x 30 µm channel in a microfluidic chip using a syringe pump at a constant flow rate of 0.16 µl/sec. For fluorescence excitation two solid-state lasers (OBIS 488-nm LS 60 mW; OBIS 561-nm LS 50 mW; Coherent Deutschland) were used and light emission was detected using two photodiode detectors (FF555-Di03, FF03-525/50; Semrock and zt 633 RDC, Chroma Technology Corp; FF01-593/46, Semrock). An image of every cell was acquired at the end of the channel using a CMOS camera (MC1362, Mikrotron) which is mounted to an inverted microscope (Axiovert, Carl Zeiss AG). In real-time cell cross-sectional area (size; $\mu m^2$) and deformation were computed. Since cell deformation and size are not independent parameter, a numerical model was to estimate corresponding Elastic modulus (6).

**Scanning electron microscopy (SEM)**

Hydrogels were immersed in SEM fixation buffer (1% glutaraldehyde, 137mM NaCl, 5mM KCl, 1.1mM $Na_2HPO_4$, 0.4mM $KH_2PO_4$, 5.5mM Glucose, 4mM $NaHCO_3$, 2mM $MgCl_2$, 2mM EDTA, 5mM MES, pH 6.5) for 1 hour. After washing in PBS, spheroids were released from the hydrogels by digesting the gel structure for 15min at 37 °C by 2.5 mg/ml collagenase (0.45 U/ml, PZ units according to Wuensch, Serva, Heidelberg, Germany). Spheroids were then kept in SEM fixation buffer overnight. Then spheroids were washed with phosphate buffer three times and stained with 1% Osmium tetroxide. Samples were dehydrated by consecutive washes in rising ethanol concentrations (50%, 70%, 90%, 95% and 100%) for 15min each. Spheroids were then transferred into porous containers and critical point dried (Leica CPD 300). Spheroids were strewn on double sided adhesive conductive tape on the SEM sample holder and sputter coated with gold (60mA, 1min, Baltec SCD 050). Samples were imaged with a table top SEM (Hitachi TM1000, Tokyo, Japan).

### Supplementary Figures

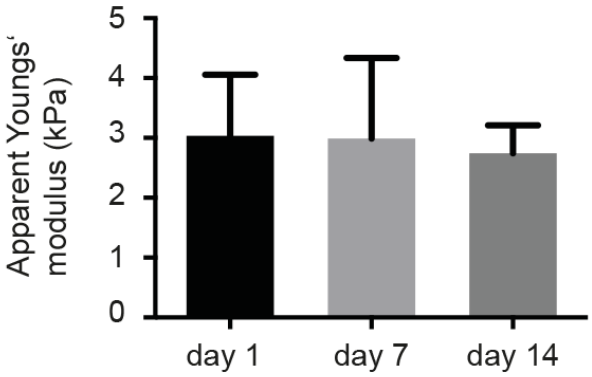


**Figure S1. Mechanical characterization of PEG-heparin hydrogels.** Hydrogels containing MCF-7 cells were probed with AFM over a period of 14 days. Data are presented as mean ± standard deviation.

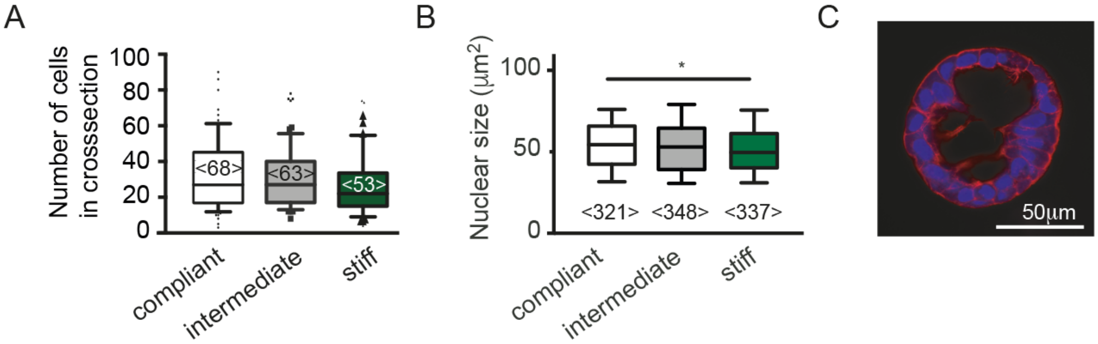


**Fig S2. Morphology of tumor spheroids in hydrogels of different hydrogel stiffness. A.** Bar graph representing the number of cells in cross sections (at the spheroid center) in confocal images of F-actin/nuclei stained 3D cultures. Data are presented as Box-Whisker plots (boxes mark the 25, 50 and 75 percentiles, whiskers mark the 10 and 90 percentiles) **B**. Nuclear cross sectional area was determined in confocal images of DAPI stained frozen sections. **C**. Example of a MCF-7 spheroid with lumen formation. The number of analyzed spheroids (A) and nuclei (B) (from at least 6 different gels per conditions) is shown in brackets. Datasets in A and B were compared using a Kruskal Wallis test and Dunns multiple comparison test. **** denotes p-values of $p<0.001$, ** $p<0.01$, * $p<0.05$.

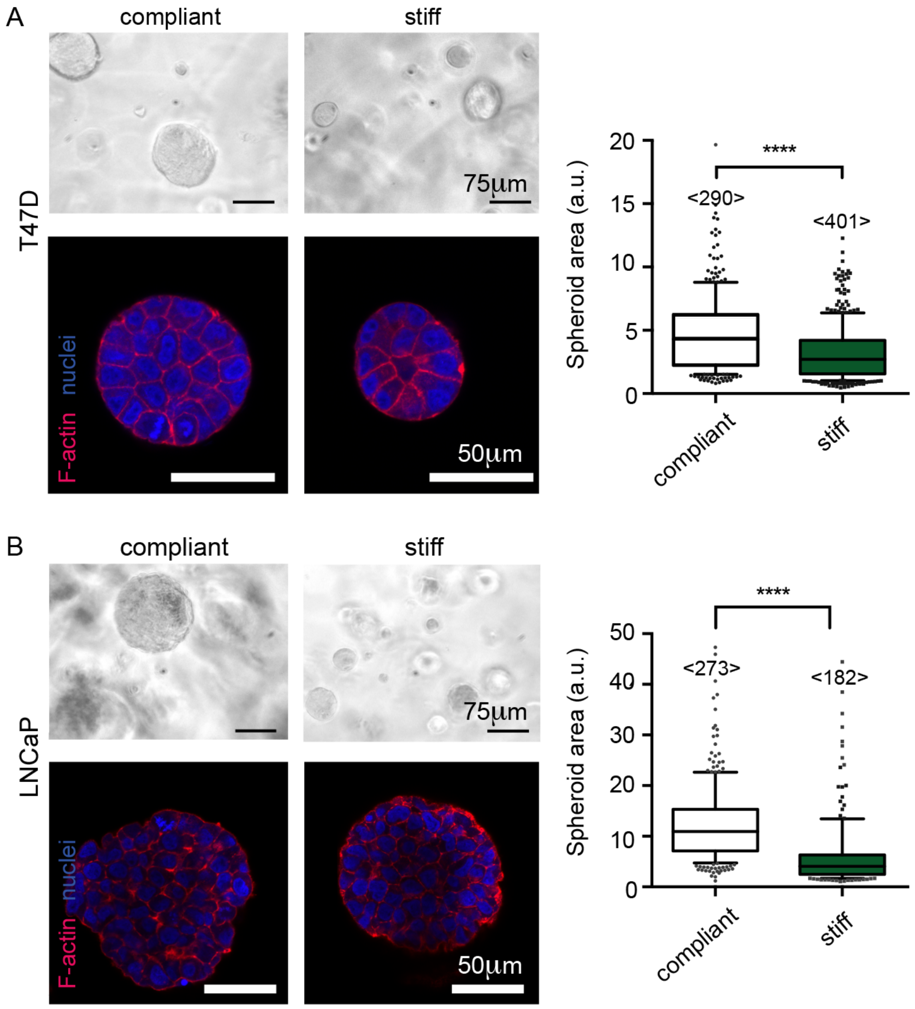


**Figure S3. Analyzing the effect of hydrogel stiffness on spheroid size in T47D and LNCaP cells.** Phase contrast (above) and confocal images of F-actin (red) and nuclei (blue) stained tumor consisting of T47D (**A**) and LNCaP (**B**) cells. The bar graphs represent spheroid diameters measured from phase contrast images of released spheroids. Data are presented as box plots (boxes mark the 25, 50 and 75 percentiles, whiskers mark the 10 and 90 percentiles). <numbers> indicate number of analyzed spheroids. Datasets were compared using a Mann-Whitney test. **** denotes p-values of < 0.001.

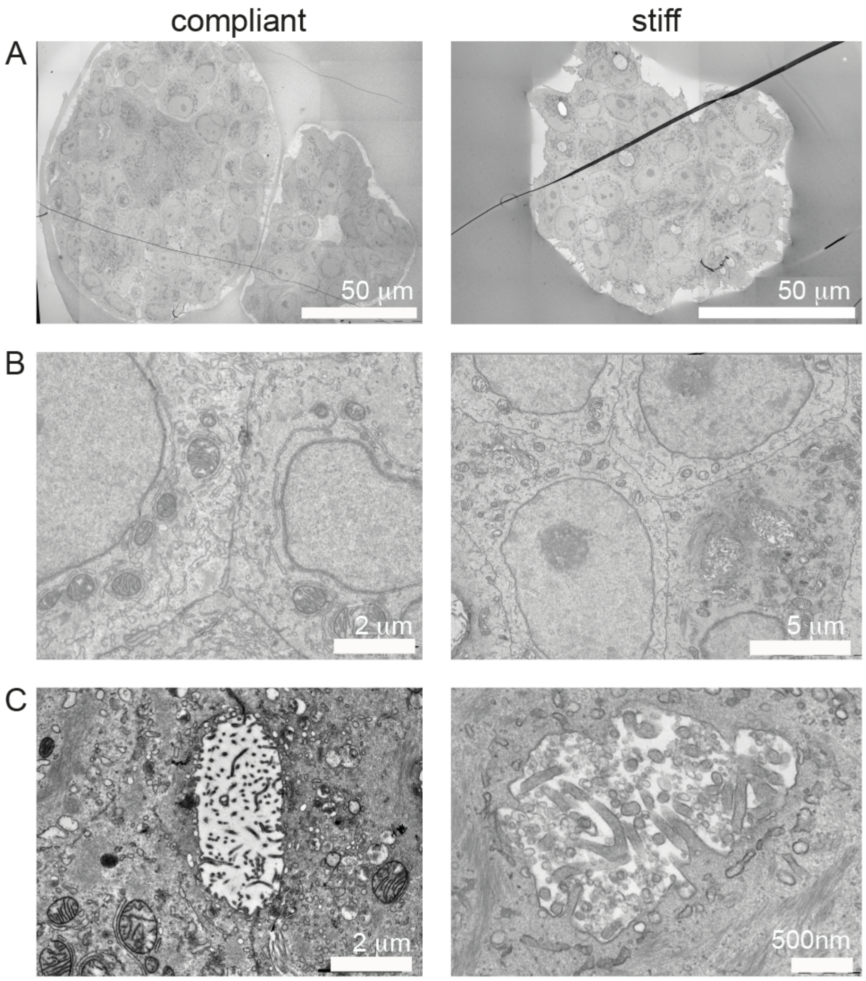


**Figure S4. TEM images of tumor spheroids in compliant and stiff hydrogels.** Overview images (**A**) and zoom-ins (**B, C**) confirm that cells are viable in compliant and stiff hydrogels. **B.** Zoom-ins highlighting examples of cell-cell contacts. **C.** Within the tumor spheroids, pores are occasionally seen, indicating secretory activity.

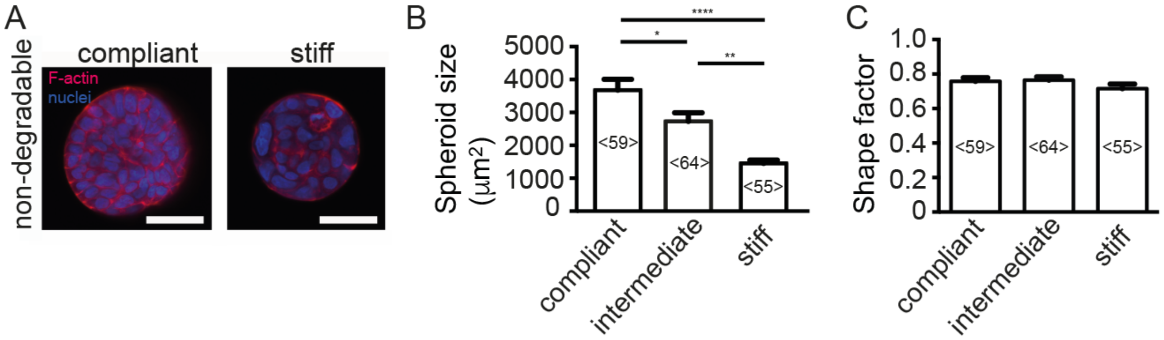


**Figure S5. Tumor spheroids grown in nondegradable hydrogels. A.** Confocal images of tumor spheroids stained for F-actin (red) and nuclei (blue). **B.** and **C.** Quantification of tumor spheroid cross sectional area (**B**) and shape factors (**C**) determined from confocal images using FIJI. Data are presented as bars showing mean ± standard error on the mean (SEM). Numbers in brackets indicate numbers of analyzed tumor spheroids. Datasets were compared using a one-way ANOVA and Tukey post hoc test. **** denotes p-values of $p<0.001$, ** $p<0.01$, * $p<0.05$.

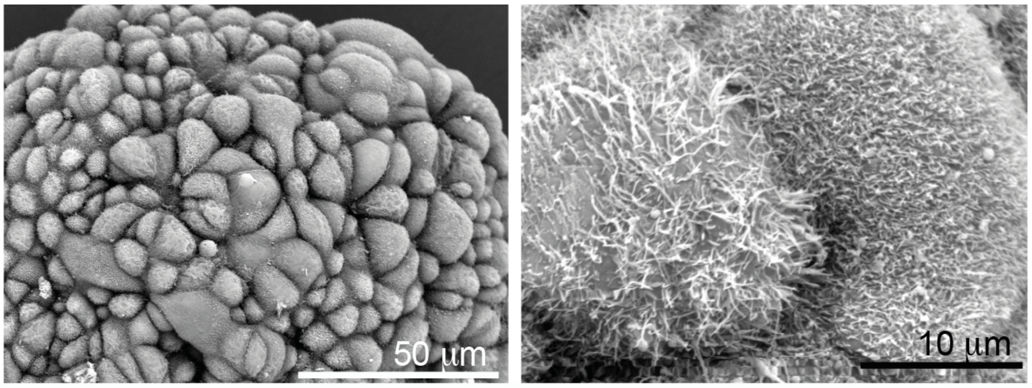


**Figure S6. SEM images of MCF-7 tumor spheroids.**

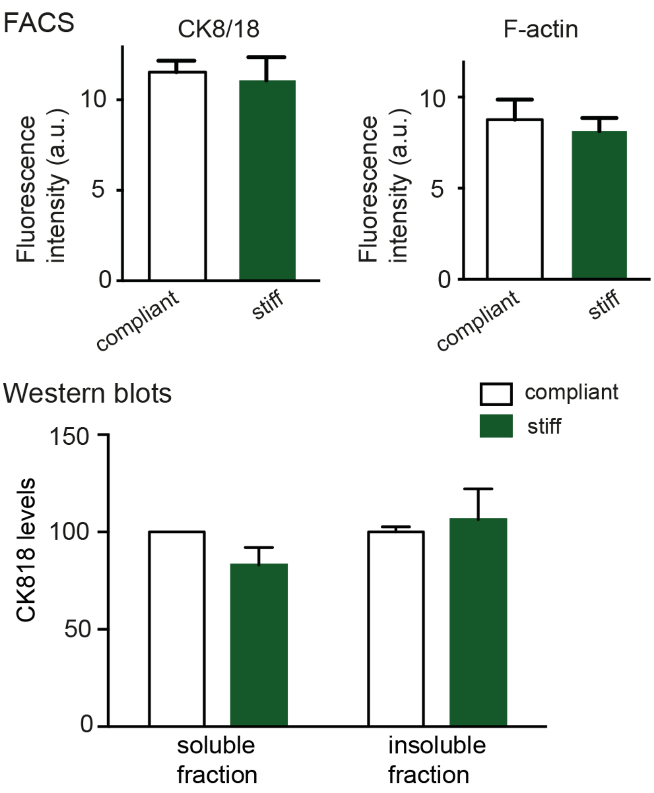


**Figure S7. FACS and Western blot data for F-actin and CK8/18. A.** FACS analysis showing total fluorescence intensity of cells stained with antibodies against CK8/18 or Phalloidin-TRITC (F-actin). **B.** Results from Western blot analysis for CK8/18 of lysates of MCF-7 3D cultures grown in compliant and stiff hydrogels (soluble and insoluble fractions). Data from 3 independent experiments were normalized to soluble factions/compliant hydrogels. Data are presented as mean ± SEM.

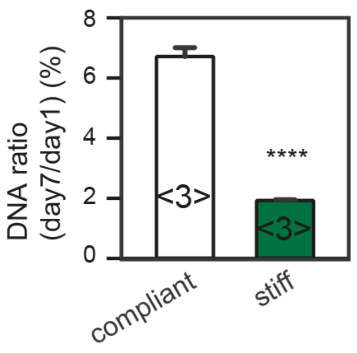


**Figure S8. Analyzing change in DNA mass of spheroid cultures in dependence of hydrogel stiffness** DNA levels were measured on day 7 and 1 using picogreen assay. The ratio of both is shown as mean ± SEM. Datasets were compared using a t- test. **** denotes p-values of $p < 0.0001$.

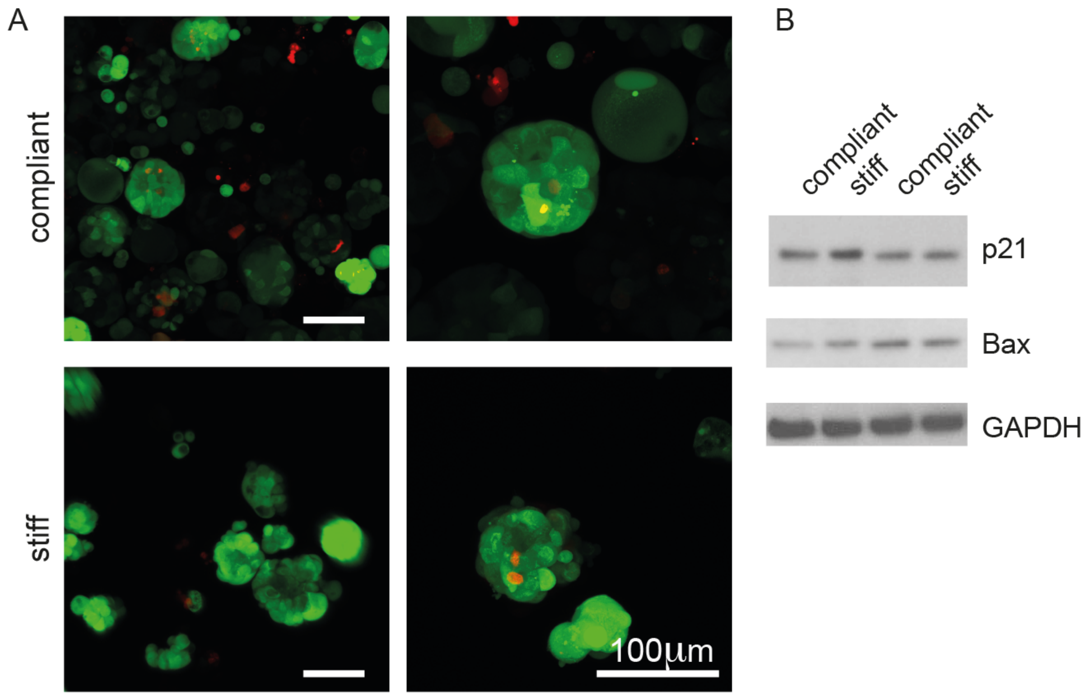


**Figure S9. Analysing cell viability in compliant and stiff hydrogel cultures. A.** 3D cultures were stained with FDA (green)/PI (red), released from gels and imaged using a confocal microscope. **B.** Western blot analysis using lysates from compliant and stiff gels for p21, Bax and GAPDH (loading control). Two independent experiments are shown.

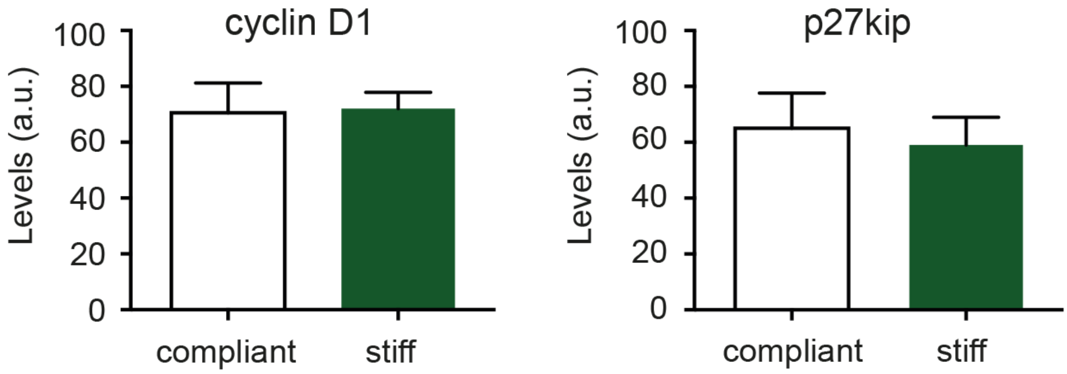


**Figure S10. Western blot analysis for cyclin D1 and p27kip.** Densitometric analysis of western blots using lysates from compliant and stiff gels for cyclin D1 and p27kip. Signals were normalized to loading controls (tubulin). Data are shown as mean ± SEM (n=3).

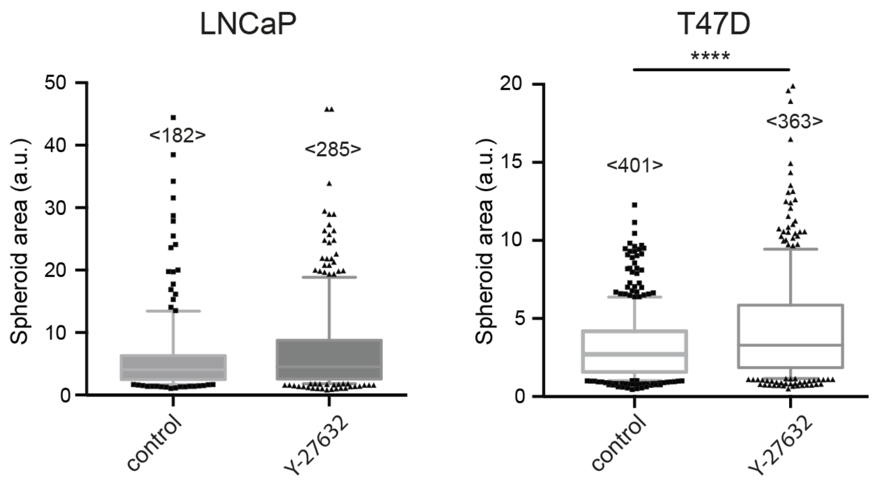


**Figure S11. Analyzing the effect of Y-27632 on T47D and LNCaP spheroid growth.** Spheroid cultures were treated from day 10 on for 96 hrs with 10uM Y-27632. After spheroid release from gels, the spheroid projected area was determined. Data are presented as Box-Whisker plot (boxes mark the 25, 50 and 75 percentiles, whiskers mark the 10 and 90 percentiles). <numbers> indicate the number of analyzed spheroids. Datasets were compared using a Mann-Whitney test. **** denotes p-values of p<0.001.

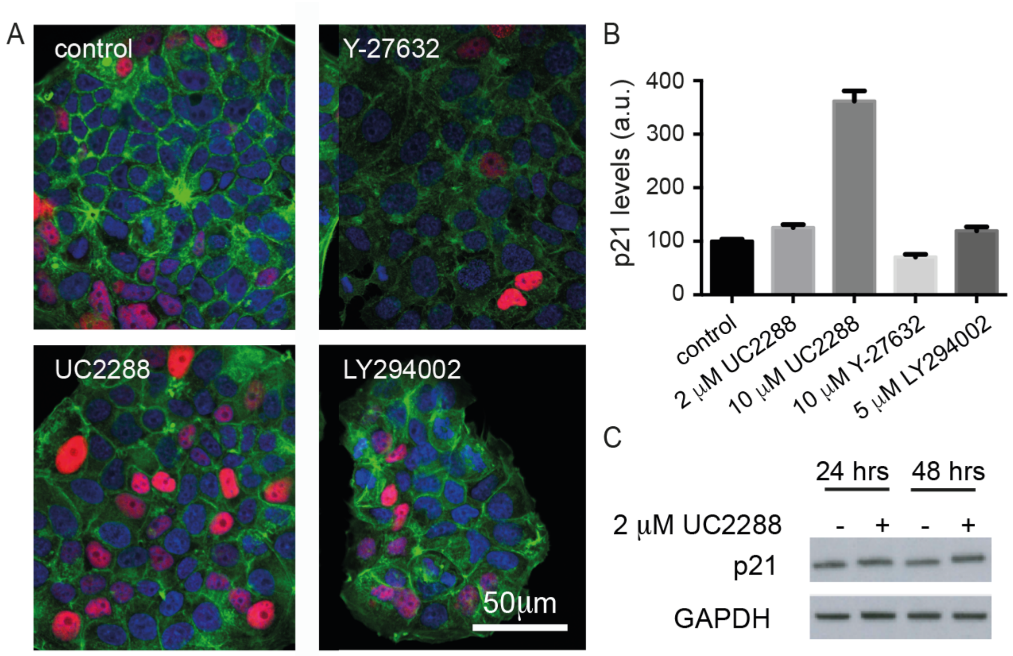


**Figure S12. Analyzing p21 levels in inhibitor treated 2D MCF-7 cultures. A.** Confocal images of MCF-7 cells stained for F-actin (green), p21 (red) and nuclei (blue) following treatment with 10 µM Y-27632, 2 µM UC2288 and 5 µM LY294002 **B.** Quantitation of staining intensities of p21 MCF-7 cells following indicated inhibitor treatment for 48 hours. **C.** Western blots of lysates from MCF-7 cells treated for 24hrs and 48hrs with or without 2 µM UC2288.

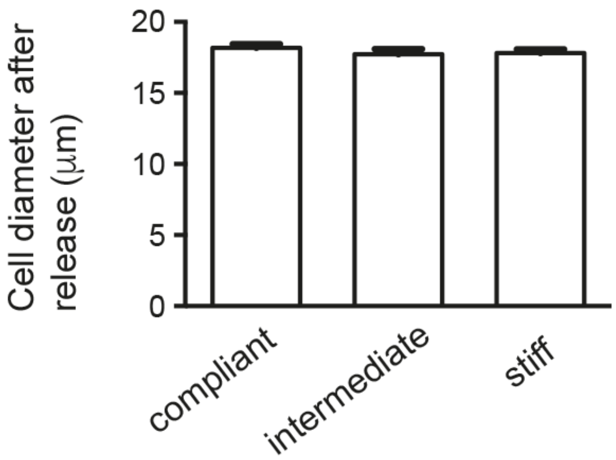


**Figure S13. Analyzing cell diameters of MCF-7 cells after release from hydrogels.** MCF-7 cells were isolated from 14-day-old spheroid cultures and imaged by phase contrast microscopy. Data are presented as mean ± SEM. A one-way ANOVA test was performed to compare datasets.

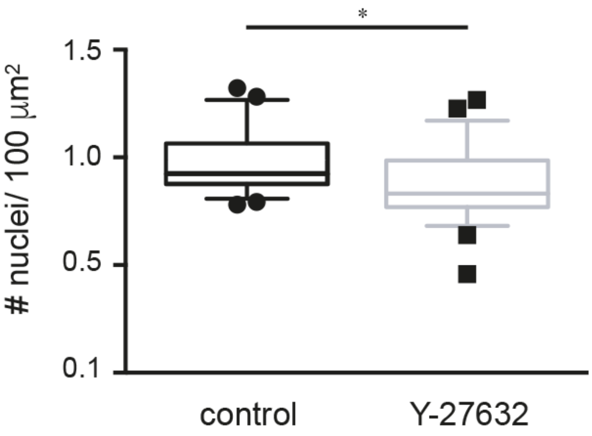


**Figure S14. Nuclear density in Y-27632 treated spheroid cultures.** Spheroids were treated from day 10 on with 10 µM Y-27632. Nuclei were counted in frozen sections stained for F-actin and nuclei. Data are presented as Box-Whisker plot (boxes mark the 25, 50 and 75 percentiles, whiskers mark the 10 and 90 percentiles). Datasets were compared using a Mann-Whitney test. * denotes p-values of $p<0.05$.

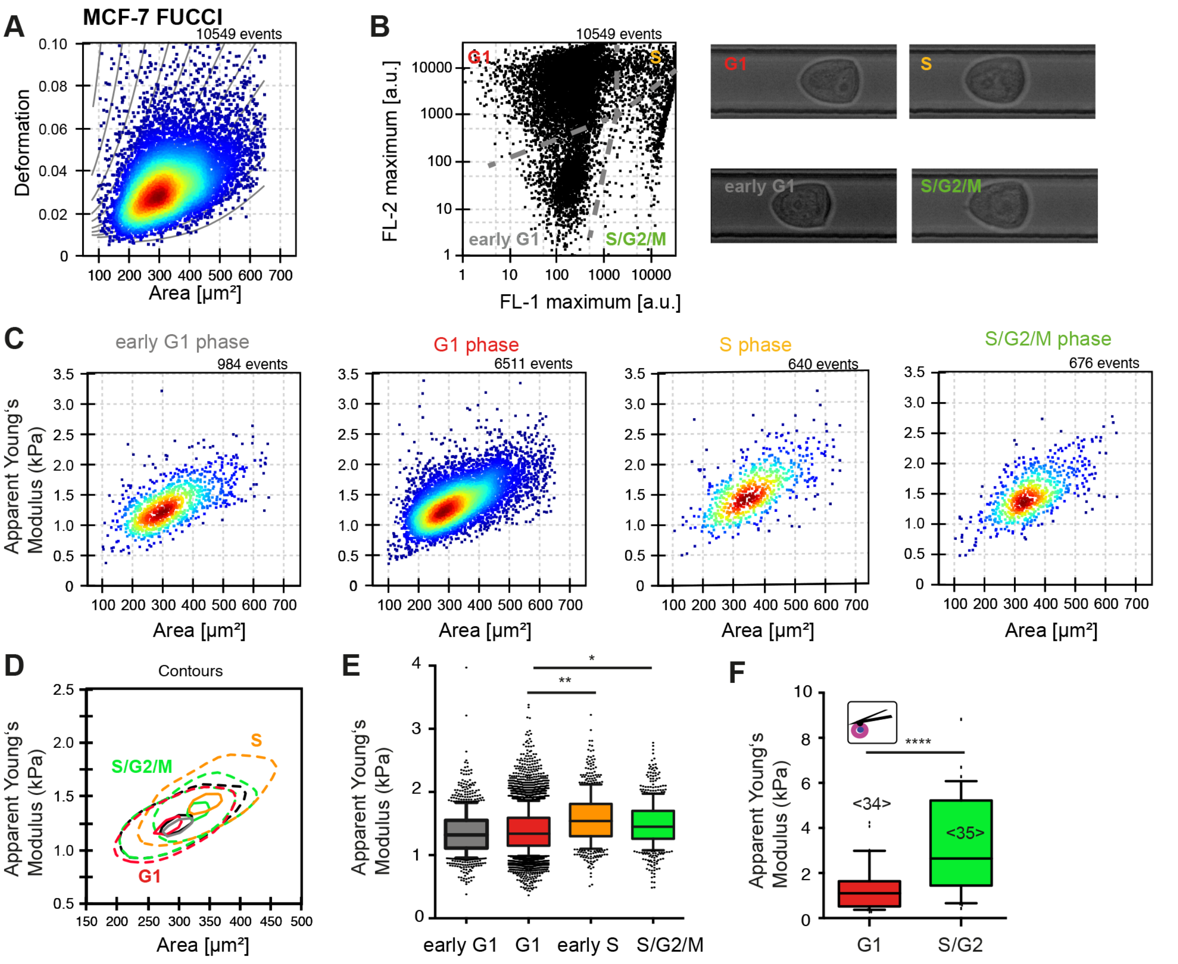


**Figure S15. Mechanical characterization of MCF-7 FUCCI cells at different cell cycle stages. A-E** MCF-7 FUCCI cells were analyzed by RTfDC. **A.** Deformation versus area plot of the whole cell population. **B.** Fluorescence channels were used to discriminate different subpopulations belonging to different cell cycle stages. Grey dotted lines indicate chosen gates. Example phase contrast images of cells in the channel are shown on the right. **C.** Apparent Young's moduli versus area plots for subpopulations defined in B. **D.** Contour plots showing apparent Young's moduli for different cell cycle stages. **E.** Box-Whisker plot of apparent Young's moduli (boxes mark the 25, 50 and 75 percentiles, whiskers mark the 10 and 90 percentiles). Indicated datasets were compared using linear mixed model analysis **F.** Apparent Young's moduli determined by AFM indentation on single MCF-7 FUCCI cells using a spherical indenter (diameter 5 µm). Datasets were compared using a Mann-Whitney test. **** denotes $p < 0.001$. <numbers> indicate the number of probed cells.
